# Dynamic transitions between neural states are associated with flexible task-switching during a memory task

**DOI:** 10.1101/2020.07.04.188235

**Authors:** Wei Liu, Nils Kohn, Guillén Fernández

**Author notes:** Correspondence: Wei Liu, School of Psychology, Central China Normal University (CCNU), No. 152 Luoyu Road, Hongshan District, Wuhan 430079, Hubei Province, Wuhan, China.

## Abstract

Flexible behavior requires switching between different task conditions. It is known that such task-switching is associated with costs in terms of slowed reaction time, reduced accuracy, or both. The neural correlates of task-switching have usually been studied by requiring participants to switch between distinct task demands that recruit different brain networks. Here, we investigated the transition of neural states underlying switching between two opposite memory-related processes (i.e., *memory retrieval and memory suppression*) in a memory task. We investigated 26 healthy participants who performed a Think/No-Think task while being in the fMRI scanner. Behaviorally, we show that it was more difficult for participants to suppress unwanted memories when a No-Think was preceded by a Think trial instead of another No-Think trial. Neurally, we demonstrate that Think-to-No-Think switches were associated with an increase in control-related and a decrease in memory-related brain activity. Neural representations of task demand, assessed by decoding accuracy, were lower immediately after task switching compared to the non-switch transitions, suggesting a switch-induced delay in the neural transition towards the required task condition. This suggestion is corroborated by an association between condition-specific representational strength and condition-specific performance in switch trials. Taken together, we provided neural evidence from the time-resolved decoding approach to support the notion that carry-over of the previous task-set activation is associated with the switching cost leading to less successful memory suppression.

**Significance statement:** Our brain can switch between multiple tasks but at the cost of less optimal performance during transition. One possible neuroscientific explanation is that the representation of the task condition is not easy to be updated immediately after switching. Thus, weak representations for the task at hand explain performance costs. To test this, we applied brain decoding approaches to human fMRI data when participants switched between successive trials of memory retrieval and suppression. We found that switching leads to a weaker representation of the current task. The remaining representation of the previous, opposite task is associated with inferior performance in the current task. Therefore, timely updating of task representations is critical for task switching in the service of flexible behaviors.

## Introduction

In everyday life, we are continuously switching between different tasks (Monsell, 2003). Transitions between task conditions have often been studied using task-switching paradigms in which participants are required to switch between two or more distinct tasks (Meiran, 2010). Usually, participants perform less accurately and/or more slowly immediately after switches (i.e., *switch costs*) (Jersild, 1927; Spector and Biederman, 1976; Rogers and Monsell, 1995; Goschke, 2000). Results from univariate fMRI studies suggested the involvement of prefrontal-parietal regions in task switching (Dove et al., 2000; Braver et al., 2003; Gruber et al., 2006; Richter and Yeung, 2014). Two studies (Waskom et al., 2014; Loose et al., 2017) used multivariate fMRI methods (Haynes, 2015; Cohen et al., 2017) to investigate how task switching modulates neural representations of the current task condition but reported mixed results. Waskom and colleagues reported stronger task representations after a shift in task conditions (Waskom et al., 2014), while Loose and colleagues found no difference in task representations between the switch and the non-switch condition (Loose et al., 2017). Therefore, it is still unclear whether task switching will strengthen or weaken the task representation and how the altered task representation links to *switch costs*.

Previous experiments that investigated task switching were typically designed to minimize the perceptual differences between conditions but not to maximize differences in underlying task demands (Braver et al., 2003; Kiesel et al., 2010; Waskom et al., 2014; Loose et al., 2017). For example, one task could be to judge the relative size of the presented object compared to a computer monitor (LARGE; e.g., *truck*; SMALL; e.g., *carrot*). The other task would be to judge whether the object is manmade (e.g., *truck*) or natural (e.g., *carrot*)(Braver et al., 2003). However, behavioral and neural correlates of task-switching between two opposite tasks within one cognitive domain remain largely unexplored. We reasoned that switching between two opposite task demands within one cognitive domain should require more cognitive resources than between unrelated tasks because the same set of networks need to reconfigure swiftly in how they interact with each other (e.g. *cooperation or competition between the very same networks*). In this study, we investigated task switching between memory retrieval and suppression and its associated neural processes in a memory task. Cognitive and neural models of memory retrieval and suppression suggest that successful retrieval could be the result of *cooperation* between an inhibitory control network and an episodic retrieval network (Rugg and Vilberg, 2013), while effective suppression depends on top-down control of the inhibitory control network upon an episodic retrieval network (i.e., *competition*) (Anderson and Hanslmayr, 2014). Previous task-fMRI studies of memory suppression supported this idea by showing that compared to Think trials, No-Think trials are associated with stronger activation in control-related regions, including the dorsolateral prefrontal cortex, ventrolateral prefrontal cortex, inferior parietal lobule, and supplementary motor area (Anderson, 2004; Guo et al., 2018; Liu et al., 2020b). At the same time, these activity increases are accompanied by reduced activity in memory-related areas in the medial temporal lobe, including the hippocampus (Anderson and Hanslmayr, 2014). A recent resting-state fMRI study showed that individual differences in memory suppression ability could be predicted by the internetwork communication of the inhibitory control network during the task-free condition (Yang et al., 2021).

Here, we used a modified Think/No-Think paradigm (Anderson and Green, 2001; Levy and Anderson, 2012) to probe task-switching between memory retrieval and suppression. Specifically, participants were instructed to switch between memory retrieval and memory suppression according to trial-specific instructions. We asked whether we can find behavioral *switch costs* (i.e., *less optimal memory performance*) when participants switch between two opposite memory tasks. If *switch costs* exist in our memory task, we would like to detect the neural source of *switch costs*. Previous cognitive theories of task-switching propose that *switch costs* could be the result of the carry-over of previous task-set activation and depends on cognitive resources required to reconfigure the task-set (Monsell, 2003). These cognitive theories can not be directly tested without the development of a series of multivariate methods to probe neural representation in non-invasive human brain imaging data (Kriegeskorte and Diedrichsen, 2019). Here, we used a time-resolved multivariate decoding approach to capture the dynamic transitions between neural states (i.e., *Think: cooperation between memory and control network; No-Think: competition between memory and control network*) during task switching in fMRI data. We hypothesized that a delayed transition between neural states that represent task conditions could be the neural underpinning of behavioral *switch costs* because failing to update neural states on time could result in a neural state that is optimal for the opposite (e.g., *retrieval*), but not the current (e.g., *suppression*) task condition. This assumption is built on the idea that the human brain can demonstrate diverse brain states during different cognitive tasks or environmental demands, and whether the brain can properly reconfigure its state is behavioral relevant (Hermans et al., 2011; Gonzalez-Castillo et al., 2015; Sadaghiani et al., 2015; Shine et al., 2016, 2019; Westphal et al., 2017; Shine and Poldrack, 2018; Cocuzza et al., 2019). The task-switching paradigm is suitable to study such a rapid neural reconfiguration process because it allows us to compare directly how different task demands are represented in neural states and how transitions of neural states are associated with human behaviors (i.e., *switch costs*).

## Results

### Behavioral results

Our study used a modified think/no-think (TNT) paradigm (**Figure 1A**) with trial-by-trial reports of (in)voluntary memory retrieval (i.e., retrieval/intrusion frequency rating) (Levy and Anderson, 2012). At the cue phase of each trial, the participant received a trial-specific instruction to either retrieve the memory that is associated with the cue (i.e., *Think trials*) or suppress the tendency to recall the memory (i.e., *No-Think trials*). Then, during the subsequent report phase, participants reported how well they just retrieved (i.e., *retrieval rating*) or suppressed the memory (i.e., *intrusion rating*). As intended, during the entire experiment (without considering the repetitions of presenting memory cues), most of the associations were successfully recalled in Think trials (1-mean p_(Never)_=84.05%, SD=11.79 %, range from 56.25% to 100%; **Figure S1A**), while participants suppressed memory retrieval successfully in No-Think trials in about half of the trials (mean p_(Never)_ =50.62%, SD=25.35%, range from 4% to 92.5%; **Figure S1B**). In addition, we investigated the learning/practicing effect by analyzing participants’ memory retrieval and suppression performance as the function of repetition: for memory retrieval trials, the percentage of reporting “always” (i.e., *index of more successful retrieval*) increased (F [9, 234] = 5.3, p < 0.001, = 0.02); for memory suppression trials, the percentage of reporting “never” (i.e., *index of more successful suppression*) increased (F [9, 234] = 5.4, p < 0.001, = 0.04) from the first to the tenth repetition. These results together suggest that participants were getting more successful at retrieving or suppressing memory traces throughout the experiment.

**Figure 1.**
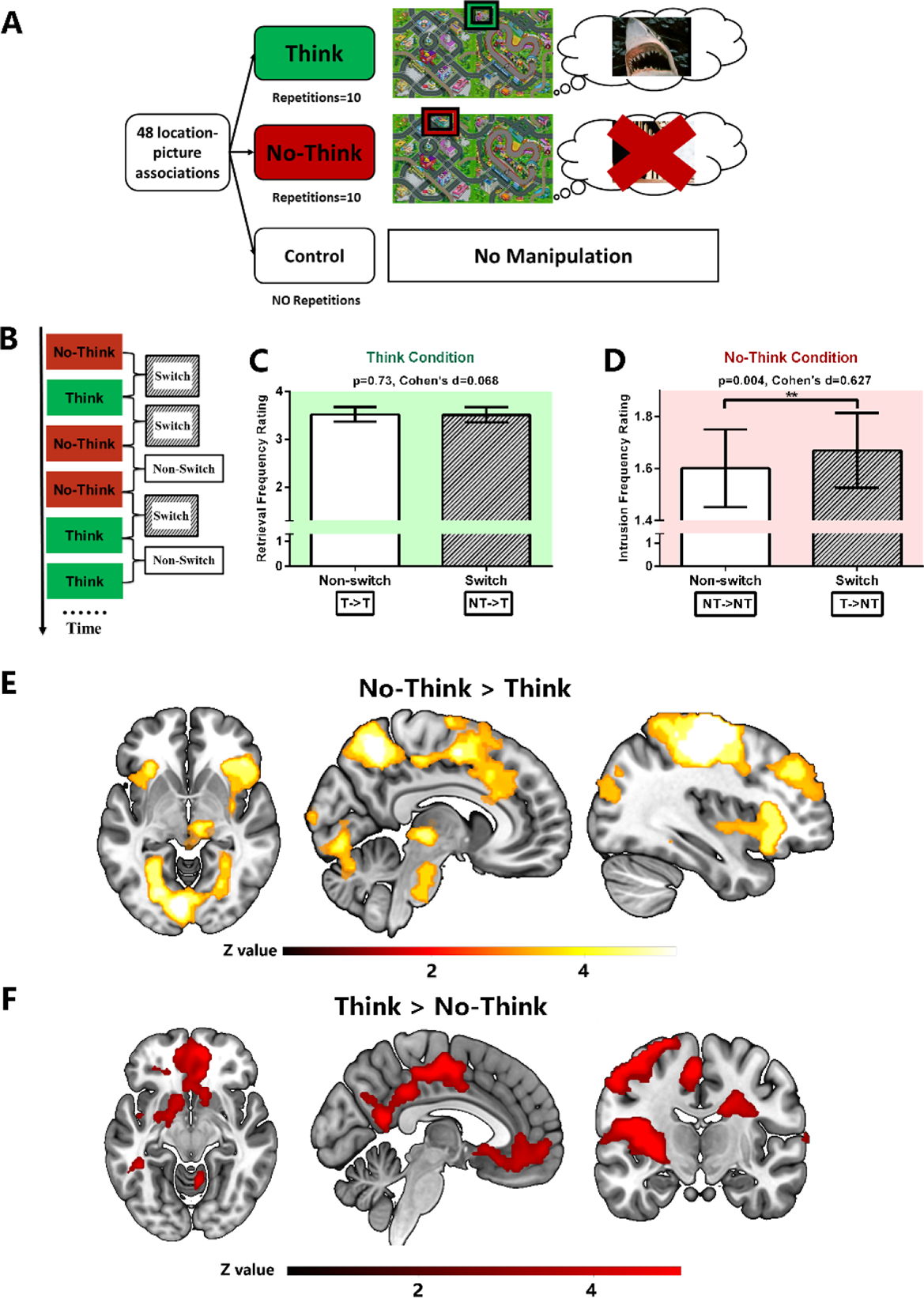
(A) After learning 48 location-picture associations, participants performed a Think/No-Think task while brain activity was measured by fMRI. During think trials, participants were instructed to retrieve associated pictures based on the highlighted locations as memory cues. By contrast, during no-think trials, participants were required to suppress the tendency to retrieval the associated pictures. **(B)** The sequence of trials was designed to probe task switching between two task demands (i.e., Think and No-Think). When the task demand of the current trial was the same as the previous trial, it was defined as the “Non-switch” trial. By contrast, while the task demand of the current differed from the previous trial, it was defined as the “switch” trial. **(C)** During Think trials, participants demonstrated comparable memory retrieval performance (p=0.73, Cohen’s d=0.068) for both “switch” (*No-Think-to-Think; NT->T*) and “Non-switch” (*Think-to-Think; T->T*) trials. **(D)** During No-Think trials, participants reported worse memory suppression perfor mance, indexed by more memory intrusions for “switch” trials (*Think-to-No-Think; T->NT*) compared to “non-switch” trials (*No-Think-to-No-Think; NT->NT*) (p=0.004, Cohen’s d=0.627). **(E)** Brain regions showed increased activation during No-Think trials compared to Think trials. **(F)** Brain regions showed increased activation during Think trials compared to No-Think trials. *Whole-brain brain imaging was thresholded at voxelwise Z>3.1, cluster-level p < .05 FWER corrected*. *For (C) and (D), Bar charts demonstrated the Mean and 95% Confidence Interval (CI) of the switch and non-switch conditions. For (E) and (F), unthresholded statistical maps can be found in the corresponding Neurovault Repository (see Data Availability) for 3D visualization*.

Our central aim was to determine whether there were behavioral *switch costs* in the TNT task and to reveal their neural underpinnings. We defined each trial as a “switch” or “non-switch” trial considering both the task condition of the current trial and its predecessor (**Figure 1B**). Specifically, we identified “switch” trials if a preceding trial had the *opposite* task condition (e.g., previous trial: Think; current trial: No-Think; T->NT). By contrast, if the current trial and the preceding trial had the *same* task condition, then the current one was a “non-switch” trial. Within each run, the number of “switch” and “non-switch” trials are almost identical (i.e., 32 vs. 31). We compared the trial-by-trial performance between “switch” and “non-switch” trials for the Think and the No-Think condition separately. Participants showed comparable performance for “switch” and “non-switch ” trials in the Think condition (t(25)=0.348, p=0.731, Cohen’s d=0.068; **Figure 1C**), while they reported more memory intrusions for “switch” trials compared to “non-switch” trials in the No-Think condition (t(25)=3.19, p=0.004, Cohen’s d=0.627; **Figure 1D**), suggesting *switch costs* when the task demand switched from a previous Think trial to a current No-Think trial (T->NT).

After the TNT task, participants performed a final memory test in which memory performance of Think trials, No-Think trials, and Control trials (i.e., associations that were learned but not presented during the TNT task) was assessed. Behavioral results from the final memory test had been reported in another publication in detail (Liu et al., 2020a) and *supplemental materials* (**Figure S2**) of this study. In this study, we used memory performance during the final memory test to quantify individual differences of *suppression-induced forgetting effects*. The *forgetting effect* was defined as the differences between memory performance between the No-Think trials and Control trials in the final memory test. More specifically, we calculated individual differences in both subjective and objective *suppression-induced forgetting effects* based on subject and objective memory measures and correlated them with fMRI measures of neural state transitions.

### fMRI results

To replicate the univariate neural signature of memory suppression reported in prior studies (Anderson, 2004; Levy and Anderson, 2012; Anderson and Hanslmayr, 2014), we first conducted a univariate analysis to contrast brain regions engaged in memory suppression and memory retrieval (i.e., *No-Think vs. Think*). We found an increased activity for No-Think trials in regions that are consistently involved in memory suppression, including the bilateral dorsolateral prefrontal cortex (DLPFC), bilateral insula, bilateral inferior parietal lobule (IPL), supplementary motor area (SMA), and middle cingulate gyrus (voxelwise Z>3.1, cluster-level p < .05 FWER corrected) (**Figure 1E**; **Table S1**). Additionally, we found higher activity in ventral visual areas and the right thalamus during No-Think compared to Think trials.

Next, we contrasted the Think condition with the No-Think condition and found the increased activity for the Think condition in a set of regions including the medial prefrontal cortex (mPFC), posterior cingulate cortex (PCC), hippocampus, inferior parietal lobule (IPL), precuneus, angular gyrus, and cerebellum (voxelwise Z>3.1, cluster-level p < .05 FWER corrected) (**Figure 1F**; **Table S2**). Together with the behavioral results from the final memory test, these results confirmed that participants in our experiment followed task instructions, leading to univariate neural signatures of memory retrieval and suppression consistent with prior findings (Anderson, 2004; Levy and Anderson, 2012; Anderson and Hanslmayr, 2014), as well as recent meta-analyses of memory suppression (Guo et al., 2018; Liu et al., 2020b).

### The transition of large-scale neural states from memory retrieval to memory suppression

Based on neurocognitive models of memory suppression (Anderson and Hanslmayr, 2014), we focused on the neural dynamics within the inhibitory control network and the memory retrieval network. First, we used *Neurosynth* (https://neurosynth.org/), an automatic meta-analysis tool of neuroimaging data (Yarkoni et al., 2011), to identify the inhibitory control network and memory retrieval network independently from our fMRI data. Using the term “*inhibitory control*” and “*memory retrieval*,” we performed term-based meta-analyses to reveal two distinct brain networks of inhibitory control (**Figure S3A**) and memory retrieval (**Figure S3B**) separately. The two meta-analytic maps have *overlapping* areas, including the IFG, insular, SMA, inferior parietal lobule (**Figure S3C**). Interestingly, the latter areas are highly similar to a “task switching” map generated by *Neurosynth* using the term “*task switching*” (**Figure S3D**).

In the next step, we tried to identify individual brain regions within the *inhibitory control*, *memory retrieval,* and *overlapping* networks. Based on the combination of a connectivity-based neocortical parcellation (number of parcels=300) (Schaefer et al., 2018) and subcortical regions (number of regions=14) (*Details see Methods*), we identified 71 regions (i.e., *memory-related regions*) within the *memory retrieval* network, 29 regions (i.e., *control-related regions*) within the *inhibitory control* network, were categorized as, and 10 regions (i.e., *overlapping regions*) within the *overlapping* network (**Figure 2A**). Finally, for each of the 110 regions, the BOLD time series were extracted from each voxel, averaged within each region, and further processed.

**Figure 2.**
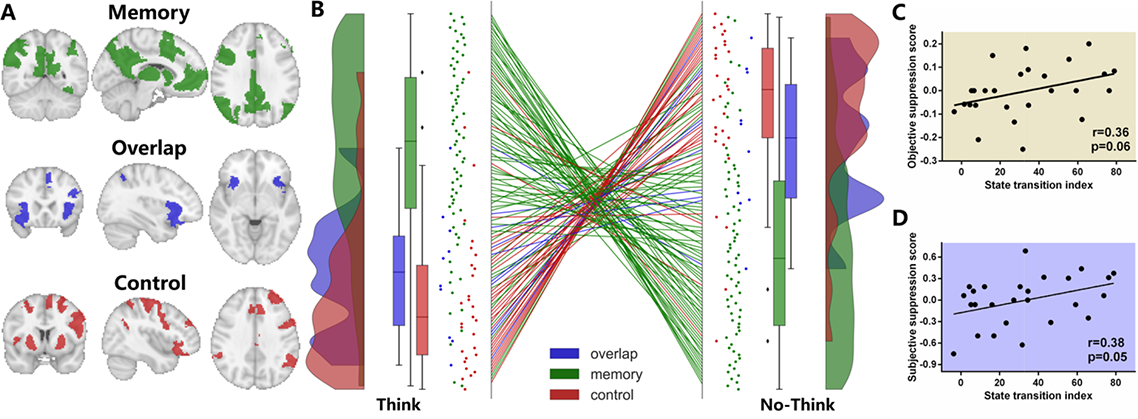
(A) Memory retrieval network (GREEN) and inhibitory control network (RED) was defined using the *Neurosynth* independent of fMRI data analyzed in this study. The overlap between the two brain networks was defined as the overlapping network (BLUE). **(B)** When the task demand switched from Think to No-Think, the activity of brain regions within the inhibitory control network increased, while the activity of brain regions within the memory retrieval network decreased. *Y-axis is the rank of neural activity among 110 regions. The top of the axis represents the highest rank (i.e., strongest activity). Each dot is a brain region.* **(C)** Individual differences in the neural state transition tended to correlate with the objective suppression-induced forgetting effect during subsequent memory retrieval task (r=0.36, p=0.06). **(D)** The same index also tended to correlate with the subjective suppression-induced forgetting effect (r=0.38, p=0.05).

Using these time series, we characterized the group-average transition of neural states when the task demand changed from Think to No-Think trials (**Figure 2B**). Based on the task instruction, the time series were firstly split for the Think and No-Think conditions separately and then concatenated across all runs of all participants. For each task demand, all regions were ranked (*the highest activity was ranked first*) based on their state-specific averaged neural activity across runs to represent their relative dominance during that neural state (i.e., *Think or No-Think*). A Kruskal-Wallis test showed that during Think trials, *memory-related regions*, *control-related regions*, and *overlapping regions* differed in their ranks (H(2)=40.48, p<0.001). Post-hoc Mann-Whitney tests using a Bonferroni-adjusted alpha level of 0.017 (0.05/3) were used to compare all group pairs. *Memory-related regions* (mean _memory_=40.22, SD _memory_=27.39) ranked higher than *control-related regions* (mean _control_=82.34, SD _control_=22.28) and *overlapping regions* (mean _overlap_=75.10, SD _overlap_=19.08) (memory-related vs. control-related: U=263, p<0.001; memory-related vs. overlapping: U=108, p<0.001). *Control-related regions* and *overlapping regions* did not differ significantly in their ranks (U=104, p=0.096). Three types of regions also differed in their ranks during No-Think trials (H(2) =36.60, p<0.001). *Memory-related regions* (mean _memory_=67.96, SD _memory_=27.94) ranked lower than *control-related regions* (mean _control_=27.24, SD _control_=23.13) and *overlapping regions* (mean _overlap_=37.80, SD _overlap_=21..10) (memory-related vs. control-related: U=238, p<0.001; memory-related vs. overlapping: U=144, p=0.001). *Control-related regions* and *overlapping regions* did not differ significantly in their ranks (U=101, p=0.081). All comparisons between *memory-related* and *control-related/overlapping regions* were significant after Bonferroni-adjustment (all ps ≤ 0.001). Furthermore, we performed additional analyses of neural state transition by dividing all regions into three groups (i.e., *increased group, stable group, and decreased group*) based on their relative changes in rank (See *Supplemental Material- Additional analyses of Think-to-NoThink neural state transition*). In short, when the task demand changed from Think to No-Think, *memory-related regions* showed decreases while *control-related regions* demonstrated an increase in their activity ranks.

For control purposes, the same analysis was repeated for raw signal intensities (**Figure S4B**) and their Z-values (**Figure S4C**) and yielded similar patterns. Here, we present parametric paired t-tests of Z-values to further validate our findings based on ranks: *control-related regions* demonstrated increased (Think>No-Think: t(28)=4.79, p<0.001, Cohen’s d=0.89) while *memory-related regions* showed decreased (No-Think>Think: t(70)=4.00, p<0.001, Cohen’s d=0.48) neural activity during the Think-to-No-Think transition. Furthermore, we analyzed the average signal within the default mode network (DMN), a functional network that is largely involved in memory retrieval, and found that DMN was less activated during the No-Think compared to the Think condition (t(25)=-3.24, p=0.003, Cohen’s d=0.63).

### The transition of neural states during the TNT task is associated with subsequent suppression-induced forgetting

We already demonstrated that the activity of *control-related regions* increased, and the activity of *memory-related regions* decreased when the task conditions switched from Think to No-Think. To assess if these changes in neural states are associated with the behavioral consequence of memory suppression (i.e., *suppression-induced forgetting effect*), we quantified this neural state transition at the individual level and examined whether individual differences in the neural state transition predict individual subsequent suppression-induced forgetting measures (i.e., *subjective and objective suppression score*). The subjective and objective *suppression score* was calculated by subtracting the memory measure (i.e., *confidence rating or recall accuracy respectively*) of suppression associations (i.e., “No-Think” items) from control associations separately.

Based on the group-level fMRI results, we calculated a *state transition index* to represent the degree of neural transition during the TNT task for each participant. The state transition index was calculated by adding up the averaged relative decreases (*absolute values for decreased values*) in ranks of all *memory-related regions* and the averaged relative increase in the rank of all *control-related regions* during the Think to No-Think transition. The larger the state transition index represents the larger decrease for *memory-related regions* and the larger increase for *control-related regions*. The *state transition index* tended to be positively correlated with individual differences in *objective suppression scores* (r=0.36, p=0.06; **Figure 2C**), and *subjective suppression scores* (r=0.38, p=0.05; **Figure 2D**). For validation purposes (*not an independent analysis*), we used an alternative method (i.e., *state transition index Version2(V2))* to measure neural state transitions for each participant. This method was based on additional analyses of Think-to-No-Think neural state transition (See *Supplemental Materials for details of state transition index calculation and results*): all regions were divided into three groups (i.e., *increased group, stable group, and decreased group*) based on their relative changes in ranks. The state transition index V2 was calculated as the sum of the percentage of *memory-related regions* within the *decreased group* and the percentage of *control-related regions* within the *increased group.* Similarly, a larger transition index V2 suggests the stronger Think-to-No-Think neural transition (i.e., *decreasing activity memory-related regions while increasing activity for control-related regions*). We also found the same significant correlations between *state transition index V2* and both *objective* and *subjective suppression scores* (**Figure S5**). These results suggested that the transition of neural states during the TNT task is relevant for the subsequent *suppression-induced forgetting effects* measured in the final memory test.

### Switch of task demand is accompanied by the delayed transition between neural states

The large-scale neural states transition described above was based on all neural data available, which means non-switch trials were also included and analyzed. To reveal how neural representations of task conditions change during task switching, we used a multivariate decoding method to track the dynamics of neural state transitions on a time point-by-time point basis (**Figure 3A**). By doing this, each time point ca be labeled as “*switch*” or “*non-switch*,” and therefore, the effect of task switching on the neural representation of task condition can be examined at the temporal resolution of each fMRI volume. Linear Support Vector Classification (SVC) was used to classify the underlying neural states (i.e., *Think vs. No-Think*) based on the fMRI activity intensity of all 110 regions at each given time point. Participant-specific classifiers were fitted on neural, and task demand data from N-1 runs (i.e., *four runs*) and tested on the one remaining test run. Then the decoding accuracy was evaluated for each TNT run by comparing the decoded task demands with the actual demands. Averaged across runs, we were able to decode task demands based on multivariate regional neural activity with a mean accuracy of 59.5% (SD=3.9% range from 52.5% to 67.1%) (**Figure 3B**). This accuracy is significantly higher than the chance level (i.e., 50%) (t(25)=12.5, p<0.001, Cohen’s d=2.453). Because we identified the learning/practicing effect at the behavioral level (i.e., *participants were getting better at retrieving/suppressing memories with repetitions*). Here, we asked whether such behavioral effects could affect the accuracies of our neural state decoding. We found that although participants’ behavioral performance improved from the first run to the last run, decoding accuracies of five runs did not differ from each other (F [4, 96] = 2.19, p= 0.075, = 0.08). Admittedly, the effect of repetition on the neural decoding is close to being significant (i.e., *decreasing, not as expected increasing, decoding accuracies throughout the experiment*). This tendency raised two things to be noted: first, because we were using the leave-one-run-out cross-validation, which assumes each run is an identical replication, the close-to-be-significant repetition effect here partly violated the assumption. Second, we suspected that the tendency of decreasing decoding accuracies may suggest that the learning/practicing process led to a less typical neural state for each task condition, and therefore allows more flexible neural state transition corresponding to the switching of external task conditions. That is the reason why we observed improved behavioral performance but a non-significant tendency of less accurate neural decoding.

**Figure 3.**
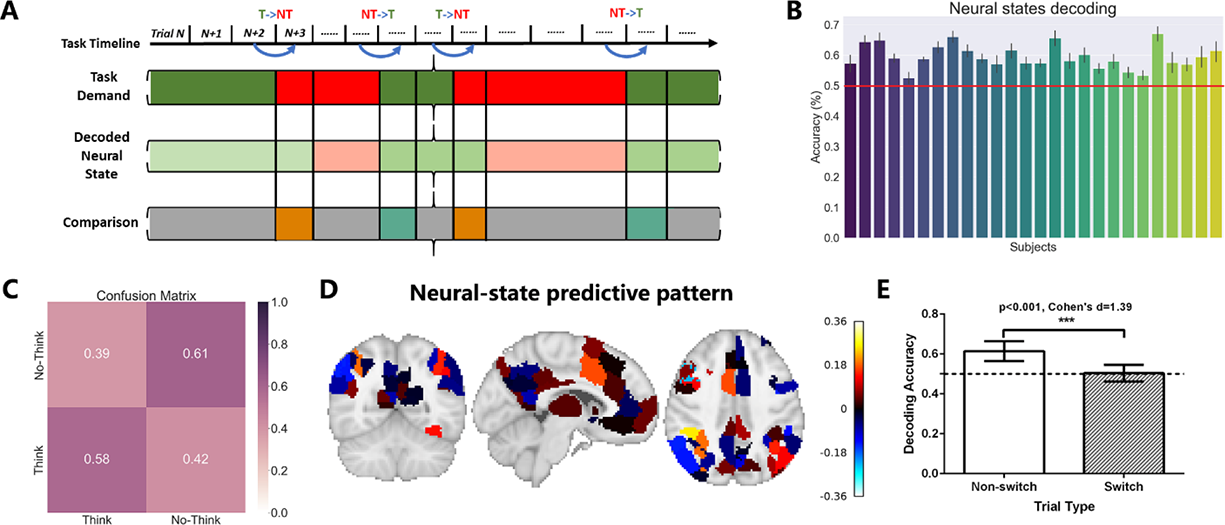
(A) Neural state decoding analysis. We trained the decoder based on large-scale brain network activity to classifier the task demand represented in the brain. We hypothesized that immediately after the switch of the task demand, the transition of the underlying neural state could be delayed. Therefore, the task demand could be misclassified as the opposite by the decoder. The real task demand was compared with the decoded neural state. The correctly decoded moments were labeled as “match” (e.g., Think as Think), while incorrectly decoded moments were defined as “mismatch” (e.g., Think as No-Think). **(B)** Decoding accuracies were presented for each participant and against the chance level of 50%. **(C)** Confusion matrix for four types of classification results. On average, 39% of the Think moments were labeled as No-Think, and 42% of the No-Think moments were regarded as Think moments by the decoder. These were the so-called “mismatch” moments depicted in Figure 3A. **(D)** The contribution of different brain regions during the decoding. This predictive pattern mainly includes the dACC, DLPFC, IFG, superior, and inferior parietal lobule. **(E)** More “mismatch” moments were found immediately after the task switching, indexed by the lower decoding accuracies during the switch compared to non-switch moments (p<0.001, Cohen’s d=1.39). *The dotted line represented the chance level (i.e., 50%) for decoding*.

We further generated the confusion matrix of our decoding analysis to quantify all types of correct and incorrect classifications (**Figure 3C**): 57.9% (SD=4.1%, range from 50.6% to 65.7%) of Think time points were correctly classified as Think. Among all No-Think time points, 61.1% (SD=3.9%, range from 53.9% to 68.4%) of them were correctly classified as No-Think. To reveal the relative contribution of each region to this decoding performance, we visualized the neural state-predictive pattern (i.e., SVC discriminating weights) in **Figure 3D**, which revealed a frontoparietal network of strong task demand representation, including the dorsal anterior cingulate cortex (dACC), DLPFC, IFG, superior, and inferior parietal lobule (**Table S3**). These regions were largely similar to the *overlapping network* (**Figure S6**).

To reveal how the switch of task conditions affected underlying neural state transitions, we calculated the decoding accuracy for “switch” and “non-switch” time points separately. Higher decoding accuracy represented a timely update of neural states according to the current task condition, thus stronger neural representation of task condition. Compared to “switch” time points, the task condition of “non-switch” time points can be decoded more accurately (t(25)=7.1, p<0.001, Cohen’s d=1.39; **Figure 3E**). That is to say, time points within No-Think trials following a Think trial were more often misclassified as Think trials compared to a No-Think trial following another No-Think trial. This pattern of results was also observed for Think trials.

### Stronger post-switch adaptive representation mitigates switch costs during the No-Think-to-Think transition

At the behavioral level, we found the *switch costs* during the Think-to-No-Think transition (i.e., T->NT), but not during the No-Think-to-Think transition (i.e., NT->T). Here, we asked whether this behavioral asymmetry can be explained neurally. Since our neural state decoding was performed on individual fMRI time points, to explore how task switching differentially affects the temporal dynamics of task representation within No-Think and Think trials, we analyzed decoding accuracies as the function of time (i.e., TR) separately for No-Think and Think trials (**Figure 4A-4B**). We found an asymmetric effect of task switching on the temporal dynamics of task representations for the Think and No-Think conditions. Specifically, a stronger post-switch adaptive task representation was selectively found after the No-Think- to-Think transition. Although the task representations are weakly represented at the beginning (TR=1: t(25)=-6.26, p<0.001, Cohen’s d=1.22) (i.e., *“delay” effects*), it was better represented halfway during the trials (i.e., *“adaptation” effects*), mainly the response phase, compared to the no-switch trials (TR=3: t(25)=5.79, p<0.001, Cohen’s d=1.13; TR=4: t(25)=2.5, p=0.019, Cohen’s d=0.49; TR=5: t(25)=2.96, p=0.007, Cohen’s d=0.58; **Figure 4A**). By contrast, for the No-Think condition, although there was also a “*delay*” effect immediately after the switch (TR=1: t(25)=-3.33, p=0.003, Cohen’s d=0.65), post-switch adaptation did not exist: task representations only differed at the beginning, and then became comparable during the response and fixation phase (**Figure 4B**).

**Figure 4.**
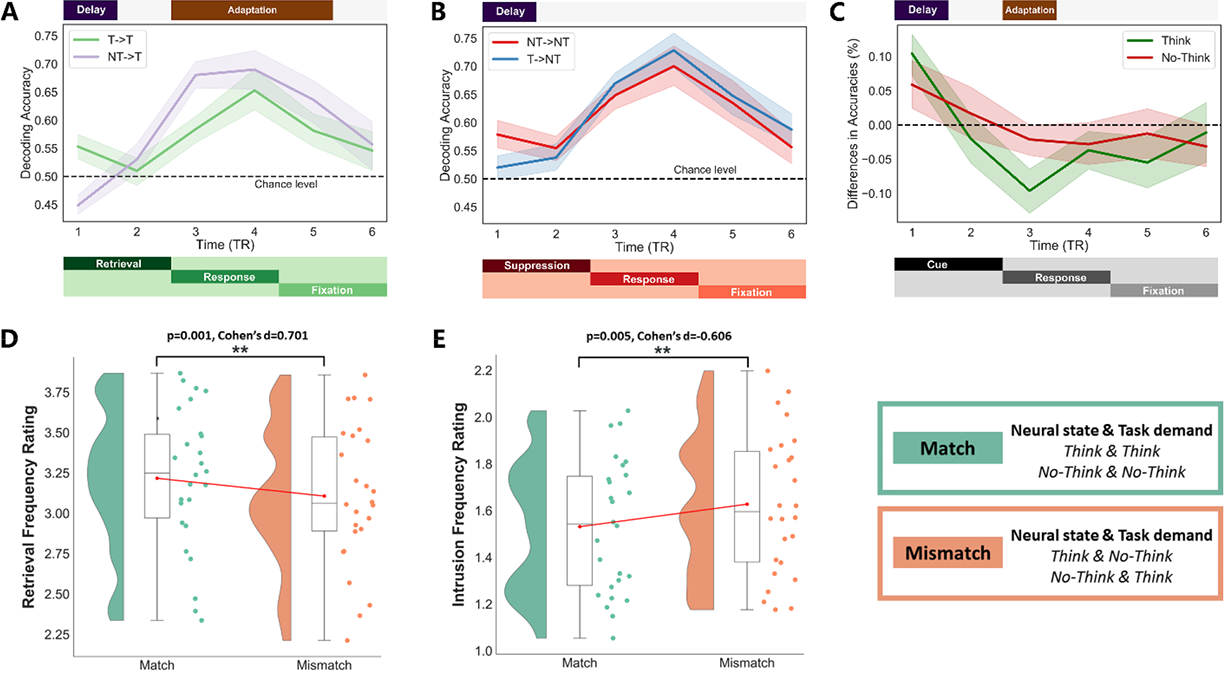
Time-revolved neural decoding of the task representation and the behavioral relevance of misclassification. **(A)** Decoding accuracies of time points during the Think condition. Lower decoding accuracies (i.e., *“delay” effects*) were found immediately after the switch (i.e., *No-Think-to-Think, NT->T*) compared to the non-switch condition, while higher decoding accuracies (i.e., *“adaptation” effects*) were found for switch compared to non-switch condition during the middle of the trials (i.e., *response phase*). **(B)** Decoding accuracies of time points during the No-Think condition. Lower decoding accuracies (i.e., *“delay” effects*) were found immediately after the switch (i.e., Think-to-No-Think, T -> NT). Decoding accuracies were comparable between switch and non-switch during the response and fixation phase. **(C)** Comparations of differences in decoding accuracies as the function of time between Think and No-Think condition. Differences were calculated by subtracting accuracies of the switch condition from the non-switch condition. Therefore, positive values (i.e., *non-swtich>switch*) represented the switch induced delay in the neural state transition, whereas negative values (i.e., *switch> non-switch*) signified the post-switch adaptive representation. *Note: this figure was made only for demonstration, the 2-by-2 ANOVA (Think/NoThink*Switch/Non-Switch) was performed to statistically test the group differences of differences.* **(D)** Memory retrieval data from Think trials. After the task switching, when the decoded neural state did not match with the task demand (i.e., *Neural state=No-Think; Task demand=Think*), participants reported worse memory retrieval performance (p<0.001, Cohen’s d=-0.70). **(E)** Memory suppression data from No-Think trials. When the neural decoder misclassified No-Think moments as Think (i.e., *Neural state=Think; Task demand=No-Think*), participants reported more memory intrusions during No-Think trials (p=0.005, Cohen’s d=0.606).

Furthermore, we calculated the differences in decoding accuracies by subtracting accuracies of the switch condition from the non-switch condition (i.e., Accuracies *_non-switch_-* Accuracies *_Switch_*) and plotted them as the function time for demonstration purposes (**Figure 4C**). To identify potential interactions, we compared these differences of accuracies between Think and No-Think trials using a 2-by-2 ANOVA (Think/NoThink×Switch/Non-Switch). There was a significant interaction effect only when the TR equals 3 (F(1,25)=12.53, p=0.002, η =0.07): decoding accuracies for the switch and non-switch condition did not differ in No-Think trials (t=-1.49, *p*_holm_=0.28), whereas decoding accuracies for the switch condition was higher than the non-switch condition in Think trials (t=6.76, *p*_holm_<0.001). To further support the idea that that higher decoding accuracies during Think trials reflect stronger task representation, and therefore, can facilitate memory retrieval, we correlated individual differences in decoding accuracies (when there were marked “*adaptation*” effects) with memory retrieval/suppression performance. We showed that individuals who demonstrated stronger “*adaptation*” effects, indexed by higher decoding accuracies, performed better at memory retrieval (*switch condition: r=0.44, p=0.02; non-switch condition: r=0.43, p=0.02*), but not memory suppression (*switch condition:r=-0.09, p=0.64; non-switch condition:r=-0.16, p-0.43*) (**Figure S7**). Although this correlation needs to be interpreted with caution and only regarded as preliminary evidence because our sample size is modest (N=26), it suggests the task-specific association between adaptive representational strength and memory retrieval.

### Mismatches between task demand and underlying neural state relate to trial-by-trial memory performance

We have already shown the association between decoding accuracies and behaviors in cross-participant analyses. Next, we sought to further link the underlying neural state to memory performance on a trial-by-trial basis. Unlike most of the decoding analyses, which usually focused on the accuracy of the classifier, here, we were particularly interested in the relationship between misclassified moments and their behavioral consequences (i.e., *switch costs).* We already demonstrated that these misclassifications were largely induced by task switching, and we predicted that this mismatch could be the neural source of behavioral *switch costs*. To test this idea, we averaged the trial-by-trial performance measures (i.e., retrieval frequency rating for Think trials and intrusion frequency rating for No-Think trials) for four situations at issue (i.e., *Think-Correct classification, Think-Incorrect classification, No-Think- Correct classification, and No-Think- Incorrect classification*) within switch trials.

We found that participants’ behavioral performance was impaired during these mismatch moments (i.e., *incorrect classifications*) (mean _incorrect_=43.9%; SD _incorrect_=2.6%; ranging from 38.5% to 49.8% of all classifications)) immediately after task switches. Specifically, during the Think condition, when neural states were mistakenly classified as No-Think, the retrieval frequency rating was lower (t(25)=-3.57, p=0.001, d=0.701; **Figure 4D**) compared to the situation in which task demands matched with the neural state. During No-Think trials, if neural states were erroneously decoded as Think, participants reported higher intrusion frequency rating (t(25)=3.08, p=0.005, d=0.606; **Figure 4E**) compared to the situation in which classifications were correct.

In our exploratory analyses, we found that such mismatch moments not only occurred during the task-switching but were also observed (*but less frequently*) during non-switch trials. Using the same decoding method but focusing on non-switch time points, we found a similar detrimental effect of mismatch on behavioral performance (**Figure S8**). These findings suggested that spontaneous, uninstructed neural state fluctuations that do not fit current task demands also have behavioral impacts.

### Applying alternative classifier training procedures did not change our findings

For the neural state decoding results presented above, we trained the classifier based on all individual fMRI time points, including not only memory-related phases (i.e., r*etrieval or suppression*) but also the response and fixation phase. This was based on our assumption that the neural representation of the current task condition remained active during the entire trial and only changed when task demands switched. One may argue that including time points beyond the mere memory-related phases (e.g., *response phase*) may bias the decoding analyses. To rule out this possibility, we re-trained our classifiers using an alternative training procedure (A) and tried to replicate our key findings. More specifically, (1) we restricted our classifier training and testing to time points during memory-related phases only (i.e., *retrieval or suppression*), (2) based on trial-by-trial memory performance, we further excluded time points when memory retrieval or suppression was unsuccessful. Classifiers that were trained based on the described alternative procedure can still decode task demands significantly higher than the chance level (t(25)=4.92, p<0.001, Cohen’s d=0.96). When we compared decoding accuracies between “switch” and “non-switch” trials, we found a switch-induced delay in neural state transitions again: lower decoding accuracy for switch trials compared to no-switch trials (t(25)=-2.42, p=0.02, Cohen’s d=0.47). Lastly, we also revealed the behavioral relevance of the misclassified time points: when the No-Think trials were misclassified as Think trials, participants reported more memory intrusions (t(25)=3.10, p=0.005, Cohen’s d=0.60); when the Think trials were mistakenly labeled as No-Think trials, participants tended to report lower rates of memory recall (t(25)=-1.98, p=0.058, Cohen’s d=0.39).

Another potential bias in our classifier training relates to the fact that we used more No-Think time points for the training than Think time points ((t(25)=4.56, p<0.001, Cohen’s d=0.89). Therefore, classifiers may be better suited to identify a “No-Think” state (i.e., *suppression state*). To control this factor, we included a subsampling balancing step before classifier training: for each participant, we randomly excluded No-Think data points to make sure that an equal number of Think and No-Think time points were used for the model training. At the same time, the original temporal sequence of the training data/label was reorganized. The latter ensures that the classifier was never aware of the actual order of the trial presentation, which should minimize the potential effects of temporal autocorrelation in the fMRI data on time-resolved decoding analyses. We re-trained classifiers and again using the described alternative training procedure (B) and replicated our three key findings: (1) classifiers can decode task condition better than chance level (t(25)=4.97, p<0.001, Cohen’s d=0.97); (2) but the decoding tended to be less accurate after task switches (t(25)=-1.83, p=0.07, Cohen’s d=0.35); (3) participants’ performances was worse during misclassified time points compared to the correctly classified time points (lower memory recall: (t(25)=-1.72, p=0.09, Cohen’s d=0.33; more memory intrusions: (t(25)=2.99, p=0.006, Cohen’s d=0.89).

Lastly, we asked whether our decoding analyses are contingent on using the specifically defined memory and inhibition networks from the Neurosynth. A recent study attempted to understand neural state transitions during task performance using whole-brain patterns of activity (Shine et al., 2019). To investigate if the delay in the neural state transitions is the network-specific or whole-brain representation, we trained our classifier based on neural activities from all 314 parcels. We found that (1) whole-brain representation can still allow us to decode task conditions at a similar level of network-specific decoding (mean _whole-brain_=59.9%, SD _whole-brain_=4.7%; ranging from 44.5% to 68.4%; mean _network-specific_=59.1%, SD _network-specific_=4.7%; ranging from 45.7% to 67.1%); (2) whole-brain representation was delayed after task switches (t(25)=-7.83, p<0.001, Cohen’s d=1.53); (3) misclassified moments were also associated with impaired memory performance (lower memory recall: (t(25)=-4.91, p<0.001, Cohen’s d=0.96; more memory intrusions: (t(25)=4.69, p<0.001, Cohen’s d=0.92)

In sum, we re-trained our decoding models to investigate whether our results are robust to three alternative classifier training procedures. Although some of the statistical tests failed to reach significance, they showed numerical trends towards our original findings, and our key decoding results can be replicated across different classifier training procedures. This outcome suggests that our conclusions are independent of specific methodological choices during classifier training.

### Neural state transitions are not the results of differences in head motion

We observed large-scale neural state transitions during the TNT task. Individual differences in these transitions were associated with the subsequent suppression-induced forgetting effect. There is a possibility that these neural state transitions are based on artifacts caused by different levels of participants’ head motion (Siegel et al., 2017; Huang et al., 2018) between Think and No-Think trials since more inhibitory control resource was required for No-Think trials compared to Think trials (Anderson and Green, 2001; Anderson and Hanslmayr, 2014). Therefore, we examined the relationship between head motion, neural state transitions, and behaviors to rule out this alternative explanation.

We analyzed the time series of head motion (i.e., *framewise displacement (FD)* (Power et al., 2012)) during the TNT task. First, there is no significant difference in mean FD values between Think and No-Think trials (FD_Think_=0.149 (SD=0.047); FD_No-Think_=0.149 (SD=0.046); t(25)=0.30, p=0.76, Cohen’s d=0.06). Second, for each participant, we calculated differences between the head motion of Think and No-Think trials (i.e., FD_Think_-FD_No-Think_) and found no correlation between these differences and *state transition indices* (r=-0.3, p=0.12)*, objective suppression scores* (r=-0.14, p=0.46) *or subjective suppression scores* (r=-0.23, p=0.25). Third, we asked whether head motion could affect our neural state decoding analysis. The head motion level tended to be lower (t(25)=-1.96, p=0.06, d=0.38) for correct decoding (FD_correct_=0.147; SD=0.045), compared to incorrect decoding (FD_incorrect_=0.151; SD=0.048). This difference raised the question of whether the lower decoding accuracy for switch compared to non-switch condition resulted from higher head motion instead of differences in the neural representation related to task demands. Therefore, we also compared head motions between the switch and the non-switch conditions. In fact, we found that head motion is even lower in the switch condition (FD_switch_=0.145 (SD=0.048); FD_non-switch_=0.150 (SD=0.047); t(25)=3.35, p=0.003, d=0.65). This result ruled the head motion out as an alternative explanation for the lower decoding accuracy for the switch condition. If the lower decoding accuracy during switching was dominantly driven by excessive head motion, we would observe relatively higher instead of lower head motion during switching. In sum, analyses of head motion suggest that our neuroimaging results are not likely to the consequence of variations in head motions between task conditions.

## Discussion

Task switching is a crucial cognitive ability that has been intensively studied using behavioral and neuroimaging methods (Meiran, 2010; Ruge et al., 2013; Richter and Yeung, 2014). Here, we investigated the task switching process between memory retrieval and suppression and demonstrated that memory suppression is more difficult when the task demand for the participants just switched from retrieval to suppression (*but not vice versa*). Applying multivariate time-resolved decoding methods to human fMRI data, we revealed that immediately after the switch, task conditions were weakly represented by the inhibitory control and memory retrieval networks, indexed by the lower decoding accuracy, compared to non-switch trials. Importantly, during the switching, when the neural representation of task demand cannot be updated in time to match the current condition (i.e., *the mismatch between task demand and neural state*), participants reported more memory intrusions in No-Think trials and less memory retrieval in Think trials. Together, we provided neural evidence based on the decoding approach to support the previously proposed cognitive theories of *switch costs*: delayed transition of task-related neural states is associated with behavioral *switch costs*. That is to say, if the neural state cannot be timely updated after the switch of task demand, behavioral performance is compromised.

In the current study, participants were instructed to perform one of two opposite memory-related tasks (i.e., *memory retrieval and memory suppression*), with the task demand staying the same or switching between consecutive trials. Similar to what was reported in the classical task-switching paradigms, which focused on reaction time and/or accuracy (Jersild, 1927; Spector and Biederman, 1976), we found *switch costs* in the memory performance. Interestingly, *switch costs* in our study were specific to memory suppression: participants reported more memory intrusions when the current No-Think trial followed a Think trial (i.e., *Think-to-No-Think transition*), suggesting a higher demand for cognitive control over the tendency to retrieve during switch trials compared to non-switch trials. At the same time, we did not find the effect of task switching on the No-Think-to-Think transition. Asymmetric *switch costs* were studied as the results of sequential difficulty effects during task switching, although the results and predictions from previous studies are mixed. Results from our study can contribute to this field if we hold the assumption that the No-Think condition is more difficult than the Think condition due to the larger executive control requirement. One previous task-switching study reported greater *switch costs* when switching from the easy to the difficult task (Arbuthnott, 2008), which is consistent with the *switch costs* only in the Think-to- No-Think transition. But some studies proposed a larger *switch cost* for the easy task than for the difficult task caused by the need for inhibition (Schneider and Anderson, 2010; Mosbacher et al., 2020). Here, we would like to acknowledge the alternative interpretation of our results: because potential differences in task difficulties could be a confound in task decoding (Todd et al., 2013), our decoding results could reflect the non-specific difficulty differences instead of memory-specific effects. Nevertheless, our time- resolved neural decoding analysis revealed the potential neural sources of these asymmetry *switch costs* (*see detailed discussion below*), and could be used to study the neural underpinnings of how difficulty leads to asymmetric *switch costs* more generally beyond memory in future studies.

The carry-over effects of the previous task-set activation on the performance of the following task have been widely studied in the task-switching literature (Monsell, 2003), but not within the context of memory. Hulbert and colleagues reported a carry-over effect of memory suppression on the subsequent memory formation (Hulbert et al., 2016): when healthy participants suppressed unwanted memories, they were more likely to fail to encode information that was presented after a suppression trial. It was proposed that memory suppression created an amnesic time window, preventing the experience within the window from being transformed into long-term memory. Evidence from fMRI supported this model by showing the reduction of hippocampal activity during memory suppression trials, and the positive correlation between individual differences in decreased hippocampal activity and the extent of memory impairment across participants (Hulbert et al., 2016).

Our finding of more memory intrusions during the No-Think trials that followed a Think trial could result from a similar mechanism: the preceding Think trials creates a time window in which the hippocampus- centered memory network remains active to support retrieval. However, if the transition of the neural state is delayed, the following No-Think trials are still located within this window, and therefore more prefrontal control resources are needed to down-regulate hippocampal activity. We tested this prediction beyond the hippocampus: large-scale neural activity of the inhibitory control and memory retrieval networks were analyzed by multivariate decoding methods to track the adaptive neural state transitions. We first characterized the transitions in neural states of memory retrieval and inhibitory control networks between Think and No-Think trials. Consistent with previous models of memory suppression (Anderson and Hanslmayr, 2014), our results showed that when the task demand switched from retrieval to suppression, memory retrieval-related regions, mainly including the hippocampus and regions of DMN, decreased their neural activity, while inhibitory control-related regions, such as dACC and LPFC, increased their activity. We also examined the relationship between individual differences in the efficiency of neural state transitions and the *suppression-induced forgetting effect* measured in the subsequent final memory test and found a positive correlation between them. There are two possible explanations for how neural state transitions are related to the effect of memory suppression: either larger or smaller state transitions are associated with the stronger suppression effect. The more intuitive explanation is that larger transitions are beneficial for suppression; however, our data suggested the opposite: participants who demonstrated less neural reconfiguration showed a stronger memory suppression effect in the following final memory test. This finding is nevertheless consistent with a previous study, which demonstrated that higher intelligence is associated with less task-related neural reconfiguration (Schultz and Cole, 2016). Our data, together with this study, may suggest that less neural reconfigurations could reflect optimization for efficient (i.e., less) state updates, reducing processing demands (Schultz and Cole, 2016). This optimal task-related neural reconfiguration could then be beneficial for memory suppression.

Recent human fMRI studies revealed task representations using multivariate decoding methods. Brain regions such as the parietal cortex, medial, and lateral PFC encode the current task demands (Bode and Haynes, 2009; Cole et al., 2011; Gilbert, 2011; Woolgar et al., 2011; Momennejad and Haynes, 2013; Waskom et al., 2014; Wisniewski et al., 2015; Etzel et al., 2016) and our study provided further support for this idea by showing that neural activity patterns of these regions largely contributed to successful discrimination between two kinds of visually highly similar trials with opposite task demands (i.e., *memory retrieval and suppression*). These identified regions have been previously associated with cognitive processing such as retrieval, maintenance, the process of rules or demands during task switching (Bunge et al., 2003; Sakai and Passingham, 2003; Gilbert, 2011; Woolgar et al., 2011; Reverberi et al., 2012). Beyond that, memory-related areas such as the hippocampus and regions within the DMN also contributed to the successful decoding in our study because the retrieval-demand and its associated neural activity significantly differed between Think and No-Think trials. However, whether these task representations can be modulated by external experimental manipulations and detected by fMRI signals is an ongoing debate. Task representations are modulated by factors including rule complexity (Woolgar et al., 2015), rewards (Etzel et al., 2016), difficulty (Wisniewski et al., 2015), and skill acquisition (Jimura et al., 2014), but not by variables such as task novelty (Cole et al., 2011), or intention (Zhang et al., 2013; Wisniewski et al., 2016).

Two studies directly investigated whether and how cognitive control processes during task switching modulate the neural representation of the current task demand. Waskom and colleagues found that task representations are enhanced after switches, indexed by the higher decoding accuracy (Waskom et al., 2014). However, they did not find evidence for behavioral switch costs in their sample; thus, the behavioral relevance of the reported stronger task representation (i.e., *higher decoding accuracy*) is unclear. Loose and colleagues did find the behavioral switch costs, but no modulation effect in the task representations (i.e., *comparable decoding accuracy between the switch and non-switch trials*) (Loose et al., 2017) and, therefore, Loose and colleagues proposed the switch-independent neural representations of the current task demand. Compared to the two studies mentioned above, our study found asymmetrical behavioral *switch costs* (i.e., *only for Think-to-No-Think transition, but not for No-Think-to-Think transition*). To investigate the potential neural source of these asymmetric *switch costs*, we further analyzed decoding accuracies as a function of time separately for memory retrieval and suppression. For both tasks, decoding accuracy was lower immediately after task switching. However, we found the post- switch adaptive representation of the current demand for memory retrieval. Specifically, although the representation of the current demand (i.e., *retrieval*) was weaker compared to the previous demand (i.e., *suppression*) immediately after switching, representations of retrieval demand were quickly increased and then they became even stronger than the previous demand half-way during trials. Interestingly, we also found preliminary correlational evidence that participants who showed stronger post-switch adaptive representation, performed better at the memory retrieval task. Therefore, we propose that this retrieval- specific adaptation may explain why there were no obvious *switch costs* during memory retrieval. Also, this interpretation would be consistent with the combination of stronger task representation and no behavioral *switch costs* reported by Waskom and colleagues (Waskom et al., 2014). Such demand-specific neural dynamics of adaptive task representation have never been reported before, potentially because our time-resolved decoding approach has not been used in the previous task-switching studies. Future studies with such a combination of time-resolved decoding approach and electroencephalogram (EEG) or magnetoencephalogram (MEG) may reveal even more details of this adaptive task representation during task switching. Critically, our neural state decoding analysis revealed the relationship between current task representation and behavioral performance on a trial-by-trial basis. Specifically, we showed that in switch trials, if the underlying neural state matched the external task demand, behavioral performance remained intact, while if the neural state was incorrectly represented, task performance was compromised. This pattern of results may further explain why higher decoding accuracy was reported together with limited behavioral switch costs in Waskom’s study (Waskom et al., 2014). As the adaptive coding hypothesis suggests (Duncan, 2001, 2010; Waskom et al., 2014), our findings demonstrated the dynamic adjustment of task representations can be tracked by large-scale neural activity from task-relevant brain networks on a trial-by-trial basis and provided direct evidence to associate delayed neural transitions with behavioral *switch costs*.

Our study has implications for a better understanding of both task switching and memory suppression. First, task switching is one of the central elements of executive control. Here, we studied task switching in the context of memory (i.e., *switch between memory retrieval and suppression*). We found that switch- induced delays in transitions of neural states can explain compromised memory performance after task switching. This principle could be tested and used to explain switch costs in other contexts: Using neural activity-based decoding approaches, we can track whether the neural state (*of the previous task*) lingers on, and probe its influences on the performance (*of the current task*). Second, memory suppression is an experimental paradigm to mimic attempts to intrusive traumatic memories. Understanding how we can better suppress intrusive traumatic memories might pave the way for new interventions and treatments for posttraumatic stress disorder (PTSD) and other affective disorders (Mary et al., 2020). However, memory suppression is usually challenging in daily life. Our results suggest that task switching and inherent switch cost may explain this challenge. Usually, there are far more moments when we try to retrieve the information instead of suppressing memories and it means that we need to switch more often from a more habitual retrieval state to a less often implemented suppression state. According to our empirical results, to minimize the negative effects of switch costs and maximize suppression-induced forgetting, potential suppression-based interventions may benefit from “long blocks” of suppression trials to prevent the carry- over effect of switching from retrieval to suppression.

*Limitation and future directions*. We used distinct locations on maps as memory cues to pair with pictures as to-be-remembered associations (i.e., *picture-picture pairs*) instead of word-word or word-picture pairs which were more frequently used in previous TNT studies (Anderson and Green, 2001; Anderson, 2004; Levy and Anderson, 2012). Our material (1) could potentially induce eye-movements as noise and (2) lead to accidental memory retrieval/suppression based on un-cued locations during visual search for memory cues. These explanations can be only investigated in future studies in which eye-tracking and neural responses are collected simultaneously. Furthermore, the fast sequential reactivation of memory traces during eye movement can be potentially detected based on EEG/MEG signals, whose temporal resolution is much higher than fMRI. However, despite these potential limitations, we have reason to think that the utilization of picture-picture pairs was not problematic because we replicated traditional TNT effects on both the behavioral and neural levels (Anderson and Green, 2001; Anderson, 2004; Levy and Anderson, 2012). We propose that using locations on maps as memory cues is important for neural state decoding analyses presented in this study, and decoding for episodic memory trace presented in another publication of our laboratory (Liu et al., 2020a) because visual and semantic processing can in this way be better controlled and more consistently compared to distinct picture/word cues. Furthermore, using maps as memory cues in the TNT paradigm opens exciting opportunities to probe how the organization principles of spatial memory (e.g., *cognitive map, cognitive graph, and grid-like representation* (Behrens et al., 2018; Bellmund et al., 2018; Peer et al., 2020)) affect the effects of retrieval practice (i.e., *Think*) and suppression (i.e., *No-Think*) on memory traces which are located at the same map. In this study, we intentionally prevented the co-localizing of the same manipulation on certain parts of the map. In future studies, the same manipulation (e.g. *No-Think*) can be assigned to specific locations on a particular map to see how memory suppression affects changes as a function of Euclidean distance and whether it can be generalized by a grid-like network. Lastly, because of the different levels of specificity in the task condition instructions (i.e., *Think instruction to recall specific memory while No-Think instruction to suppress all related memories and thoughts*), the Think vs. No-Think contrast is not optimal and related overall BOLD activity levels and behaviors are difficult to compare. For the neuroimaging analysis, to replicate previous memory suppression findings, we have no choice but to contrast Think trials and No- Think trials. For the behavioral data, we intentionally analyze retrieval and intrusion rating separately instead of performing the interaction analysis like the conventional task-switching studies.

In summary, our results provide neural insights into the flexible task switching between memory retrieval and memory suppression. We found evidence for *switch costs* in memory suppression: it is more difficult to suppress unwanted memories immediately after memory retrieval. During switching between retrieval and suppression, we observed delayed transitions of neural states that each of them separately represents current task demand. Delayed neural transitions were associated with *switch costs* (i.e., *unsuccessful suppression and retrieval*), which directly support previously proposed cognitive theories that *switch costs* could be the result of the carry-over of previous task-set activation. These results provide insight into the critical role of dynamically adjusted neural reconfigurations in supporting flexible memory suppression and the broader neural mechanisms by which humans can flexibly adjust their behavior in ever-changing environments.

## Materials and Methods

### Participants

In total, thirty-two right-handed, healthy young participants recruited from the Radboud Research Participation System finished all of the experimental procedures. All of them are native Dutch speakers. Six participants were excluded from data analyses due to low memory performance (i.e., lower the chance level (25%) during the final memory test) (n=2), or excessive head motion (n=4). We used the motion outlier detection program within the FSL (i.e., FSLMotionOutliers) to detect timepoints with large motion (threshold=0.9). There are at least 20 spikes detected in these excluded participants with the largest displacement ranging from 2.6 to 4.3, while participants included had less than ten spikes. Finally, 26 participants (15 females, age=19-30, mean=23.51, SD=3.30) were included in the behavioral and neuroimaging analysis reported in this study. Due to the reconstruction error during the data acquisition, one run of one participant is not complete (20-30 images were missing). Therefore, that run was not included in our analysis of time series. But unaffected acquired images of that run were used in our univariate activation analysis. No participants reported any neurological and psychiatric disorders. We further used the Dutch-version of the Beck Depression Inventory (BDI) (Roelofs et al., 2013) and State-Trait Anxiety Inventory (STAI) (van der Bij et al., 2003) to measure the participants’ depression and anxiety level during scanning days. No participant showed a sign of emotional problems (i.e., their BDI and STAI scores are within the normal range). The experiment was approved by and conducted in accordance with requirements of the local ethics committee (Commissie Mensgebonden Onderzoek region Arnhem-Nijmegen, The Netherlands) and the declaration of Helsinki, including the requirement of written informed consent from each participant before the beginning of the experiment. Each participant got 10 euros/hour for their participating.

### Experiment design

This experiment is a two-day fMRI study, with 24 hours delay between two sessions (**Figure S8**). fMRI data of the day2 final memory test has been published in another publication (Liu et al., 2020a), and the comprehensive reports of the experimental materials and design can be found there. Because all of the behavioral and neuroimaging data included in this study came from the Day2 session, we just presented a brief description of the Day1 session. On day1, we instructed participants to memorize a series of sequentially presented location-picture associations, for which 48 distinct photographs were presented together with 48 specific locations on two cartoon maps. In our experiment, we used picture (i.e. *location*)-picture pairs as to-be-remembered materials instead of word-word or word-picture pairs to keep visual processes during scanned tasks largely consistent. More specifically, whole maps were presented with sequentially highlighting specific locations by colored frames as memory cues, therefore, at the perceptual level, participants were always processing the same two maps. All photographs can be assigned into one of the four categories, including animal, human, scene (e.g., train station), and object (e.g., pen and notebooks). Therefore, objective memory performance could be assessed within the scanner by instructing participants to indicate the picture’s category when cued by the map location. During this study phase, each location-picture association was presented twice, and the learning was confirmed by two typing tests outside the scanner. During the typing tests, participants were required to describe the photograph associated with the memory cue in one or two sentences. Immediately after the study phase (Day1), 88.01% of the associated pictures were described correctly (SD= 10.87%; range from 52% to 100%).

On Day2, participants first performed the second typing test, and still recalled 82.15% of all associations (SD = 13.87%; range from 50% to 100%). Then, they performed the Think/No-Think (TNT) task, and final memory test insider the scanner. We used the TNT task with trial-by-trial performance rating to monitor the retrieval or suppression of each trial. Compared to the original TNT task (Anderson, 2004), the additional self-report did not affect the underlying memory suppression process and also was used in a neuroimaging experiment before (Levy and Anderson, 2012). Forty-eight picture-location associations were divided into three conditions (i.e., “think or retrieval,” “no-think or suppression,” and “baseline or control” condition) in a counterbalanced way, therefore, for each association, the possibility of belonging to one of the three conditions is equal. For each map, 24 locations that were distributed evenly across the map were paired with six pictures from each category. One-third of associations (8 associations; 2 pictures from each category) on that map were retrieval associations (i.e. “*Think” associations*), one-third of associations were suppression associations (i.e., *“No-Think ”associations*), and the remaining one-third were control associations. The spatial distribution of the three conditions was determined manually by the experimenter to prevent clusters of specific conditions on certain parts of the maps. During the retrieval condition, locations were highlighted with the GREEN frame for 3s, and participants were instructed to recall the associated picture quickly and actively and to keep it in mind until the map disappeared from the screen. By contrast, during the suppression condition, locations were highlighted with the RED frame for 3s, and our instruction for participants was to prevent the potential memory retrieval and try to keep an empty mind. We gave additional instructions for the suppression condition: “*when you see a location, highlighted with a RED frame, you should NOT think about the associated picture. Instead, you should try to keep an empty mind during this stage. It is a difficult task, and it is totally fine that sometimes you still think about the associated picture. But please do NOT close your eyes, focus on something outside the screen, or think about something else in your life. These strategies, although useful, could negatively affect the brain activity that we are interested in ……*”. After each trial, participants had a maximum 3s to press the button on the response box to indicate whether and how often the associated picture entered their mind during Think or No-Think trials. Specifically, they rated their experience from 1-4 representing from No Recall (i.e., Never) to Always Recall. Responses during Think trials were used as retrieval frequency ratings, while responses during No-Think trials were regarded as intrusion frequency ratings. Associations that belong to the control condition (16 associations) were not presented during this phase. The TNT task included five functional runs, with 32 retrieval trials and 32 suppression trials per run. All “retrieval” or “suppression” associations were presented twice within one run, but not next to each other. Therefore, they were presented ten times during the entire TNT task. Between each trial, fixation was presented for 1- 4s (mean=2s, exponential model) as the inter-trial intervals (ITI).

To investigate the task switching within the TNT task, for each run of each participant, we predefined the sequence of task demand to form “blocks” of memory retrieval or suppression with the length range from 1 trial to 4 trials (mean=1.9 trials, std=1.01 trials, P _one-trial block_=46.875%, P _two-trials block_=25%, P _three-trials block_=18.75%, P _four-trials block_=9.375%). In this sequence, the task demand of the current trial can be the same as the previous trial (“non-switch” trial) or differ from the previous trial (“switch” trial). Within one run of a total of 64 trials, 31 trials were “non-switch” trials, 32 trials were “switch” trials, and the first trial cannot be labeled as “non-switch” trials or “switch” trials because it has no predecessor. The “non-switch” trials and “switch” trials both accounted for around 50% of the “retrieval” and “suppression” trials. After determining the sequence of task demand, specific location-picture associations from retrieval or suppression condition were randomly selected for each trial.

After the TNT task, a final memory test was performed by participants within the scanner to evaluate the effect of different modulations on memory. All 48 memory cues (i.e., locations), including retrieval, suppression and control conditions, were presented again with the duration of 4s by highlighting a certain part of the map with a BLUE frame. Participants were instructed to recall the associated picture as vividly as possible during the presentation and then give the responses on two multiple-choice questions within 7s (3.5s for each question). The first one is the measure of subjective memory: “how confident are you about the retrieval?”. Participants had to rate from 1 to 4 representing “Cannot recall, low confident, middle confident and high confident” separately. The second one is the measure of objective memory: “Please indicate the category of the picture you were recalling.” They needed to choose from four categories (i.e., Animal, Human, Scene, and Object). It is notable that we only analyzed the behavioral data from this within-scanner memory test; the neural activity during this test is not the focus of this study.

### Behavioral data analysis

Behavioral results of this project were comprehensively reported in another study of our lab with the focus on the final memory test (Liu et al., 2020a). No results of tasks sneurvawitching (i.e., *switch costs*) were reported in that study, and task switching is the central scientific question of this study. First, we analyzed the behavioral performance during the TNT task. Trial-by-trial performance reports from each participant were used to calculate the percentage of successful recall chosen across 160 retrieval trials and successful suppression across 160 suppression trials. Following previous studies (Levy and Anderson, 2012; Liu et al., 2020a), performance reports from suppression trials were used to quantify individual differences in memory suppression efficiency (“*intrusion slope score*”). To account for the individual differences in memory performance before the TNT, we restricted the analysis of suppression into the associations for which participants can still remember during the second typing test (“remembered associations”). We used linear regression to model the relationship between intrusion frequency ratings of “remembered associations” and the number of repetitions of suppression at the individual level. Participants with more negative slope scores are better at downregulating memory intrusions than those with less negative slope scores. Furthermore, we labeled each trial as the “non-switch” trial or “switch” trial based on whether the task demand of the current trial is the same as the previous trial. Trial-by-trial performance between “switch” and “non-switch” trials during retrieval or suppression was compared using paired t-tests.

We also quantified the individual differences in *suppression-induced forgetting effect* based on two types of participants’ performance (i.e., recall accuracy and confidence rating) during the final memory test. Memory performance for associaitons that belong to the control condition was regarded as the baseline to qualtify the suppression-induced forgetting effect. For each participant, recall accuracy (objective memory measure) and confidence rating (subjective memory measure) were calculated for No-Think associations and control associations separately. Then objective and subjective *suppression scores* were computed separately by subtracting the accuracy and confidence of No-Think associations from the control associations. The more negative a *suppression score* is, the stronger the *suppression-induced forgetting effect* is. The memory suppression score was used to correlate with the “*intrusion slope score*” and transition of neural states during the TNT.

### MRI data acquisition and preprocessing

We used a 3.0 T Siemens PrismaFit scanner (Siemens Medical, Erlangen, Germany) and a 32 channel head coil system at the Donders Institute, Centre for Cognitive Neuroimaging in Nijmegen, the Netherlands to acquire MRI data. For each participant, MRI data were acquired on two MRI sessions (around 1 hour for each session) with 24 hours’ interval. In this study, we only used the data from the day2 session. Specifically, we acquired a 3D magnetization-prepared rapid gradient echo (MPRAGE) anatomical T1-weighted scan for the registration purpose with the following parameters: 1 mm isotropic, TE = 3.03 ms, TR = 2300 ms, flip angle = 8 deg, FOV = 256 × 256 × 256 mm. All functional runs were acquired with Echo-planar imaging (EPI)-based multi-band sequence (acceleration factor=4) with the following parameters: 68 slices (multi-slice mode, interleaved), voxel size 2 mm isotropic, TR = 1500 ms, TE = 39 ms, flip angle =75 deg, FOV = 210 × 210 × 210 mm. In addition, to correct for distortions, magnitude and phase images were also collected (voxel size of 2 × 2 × 2 mm, TR = 1,020 ms, TE = 12 ms, flip angle = 90 deg).

We used the FEAT (FMRI Expert Analysis Tool) Version 6.00, part of FSL (FMRIB’s Software Library, www.fmrib.ox.ac.uk/fsl) (Jenkinson et al., 2012) together with Automatic Removal of Motion Artifacts (ICA-AROMA) (Pruim et al., 2015) to perform our preprocessing. This pipeline was based on procedures suggested by Mumford and colleagues (http://mumfordbrainstats.tumblr.com) and the article that introduced the ICA-AROMA (Pruim et al., 2015). Specifically, we first removed the first four volumes of each run from the 4D sequences for the stabilization of the scanner and then applied the following pre-statistics processing: (1) motion correction using MCFLIRT (Jenkinson et al., 2002); (2) field inhomogeneities were corrected using B0 Unwarping in FEAT; (3) non-brain removal using BET (Smith, 2002); (4) grand-mean intensity normalization of the entire 4D dataset by a single multiplicative factor; (5) spatial smoothing (6mm kernel). ICA-AROMA was used to further remove motion-related spurious noise. We chose to conduct “non-aggressive denoising” and applied highpass temporal filtering (Gaussian-weighted least-squares straight-line fitting with sigma=50.0s) before the following analyses.

All of the mentioned preprocessing steps were performed in native space. We used the following steps to perform the registration between native space, participant’s high-resolution T1 space, and standard space. Firstly, we used the Boundary Based Registration (BBR) (Greve and Fischl, 2009) to register functional data to the participant’s high-resolution structural image. Next, registration of high resolution structural to standard space was carried out using FLIRT (Jenkinson and Smith, 2001; Jenkinson et al., 2002) and was then further refined using FNIRT nonlinear registration (Andersson et al., 2007). Resulting parameters were used to align processed functional images from native-space to standard space for the following signal extraction.

### Univariate General Linear Model (GLM) analyses

We ran the voxel-wise GLM analyses of the TNT task to identify brain regions that are more active during memory suppression compared to memory retrieval (i.e., No-Think VS Think). All time-series statistical analysis was carried out using FILM with local autocorrelation correction (Woolrich et al., 2001) using FEAT. In total, three regressors were included in the model. We modeled the presentation of memory cues (locations) as two kinds of regressors (duration=2TR)(i.e., suppression trials and retrieval trials). For each participant, based on the second typing test which was immediately before the TNT phase, we separately modeled the location-picture associations, which participants cannot describe (i.e., *unlearned associations*) as a separate regressor. Because these location-picture associations were unlearned/forgotten, there were no memories to be recalled or suppressed during the TNT phase. We conducted the two contrasts-of-interest (i.e., No-Think VS Think and Think VS No-Think) first at the native space and then aligned resulting statistical maps to MNI space using the parameters from the registration. These aligned maps were first used for participant-level averaging across five TNT runs, and then the group-level analyses. The group-level statistical map was corrected for multiple comparisons using default cluster-level correction within FEAT (voxelwise Z>3.1, cluster-level p < .05 FWER corrected).

### Networks-of-interest identification

To identify our networks-of-interest (i.e., inhibitory control network and memory retrieval network), we performed several term-based meta-analyses using the *Neurosynth* (https://neurosynth.org/) (Yarkoni et al., 2011). “Inhibitory control” and “memory retrieval” were used as terms separately to search for all studies in the *Neurosynth* database whose abstracts include the input term at least once. Then, all identified studies were combined separately for each term to generate the corresponding statistical map. We used uniformity test maps in our study. This method tested whether the proportion of studies that report activation at a given voxel differs from the rate that would be expected if activations were uniformly distributed throughout the grey matter. Voxel-wise Z-score from the one-way ANOVA testing was saved in a statistical map. Each map was thresholded to correct for multiple comparisons using a false discovery rate (FDR)(p<0.01). It is notable that due to the continuous update of the *Neurosynth* database, the number of studies included in the analyses could be slightly different for each search, the maps we used can be found in our *Neurovault* repository (https://identifiers.org/neurovault.collection:7731). Similar network identification was also performed using *BrainMap* (Laird et al., 2005) as a confirmation. The two methods of meta-analysis yielded highly similar maps of network-of-interests (**Figure S9**), and we used the maps generated by the *Neurosynth* in our main text.

We used the thresholded (p_FDR_<0.01) spatial maps of “inhibitory control” and “memory retrieval” to general three masks of networks-of-interest. The areas which belong to both the “inhibitory control” and “memory retrieval” masks were labeled as *overlap network*, the areas which only belong to the “inhibitory control” mask were labeled as *inhibitory control network*, and the areas which only belong to the “memory retrieval” masks were labeled as *memory retrieval network*.

### Brain parcels for the extraction of time series

We combined a parcellation of cerebral regions (N=300; mean size=440.1 voxels (SD=188.9 voxels)) (Schaefer et al., 2018) and all subcortical regions (N=14; mean size=491.57 voxels (SD=334.7 voxels) from the probabilistic Harvard-Oxford Subcortical Structural Atlas (Desikan et al., 2006) as a whole-brain parcellation. The parcellation of cerebral regions was based on a gradient-weighted Markov Random Field (gwMRF) model, which integrated local gradient and global similarity approaches (Schaefer et al., 2018). Based on both task fMRI and resting-state fMRI acquired from 1489 participants, parcels with functional and connectional homogeneity within the cerebral cortex were generated. Each parcel is one of the seven large-scale functional brain networks, including *Visual, Somatomotor, Dorsal Attention, Ventral Attention, Limbic, Frontoparietal, Default network* (Yeo et al., 2011). Subcortical regions included bilateral thalamus, caudate, putamen, globus pallidus, hippocampus, amygdala, and ventral striatum. Details of each parcel (e.g., name, coordinates, hemisphere) within the whole brain parcellation can be found in our OSF folder (https://osf.io/cq96h/). Some may argue that the size of parcels differs across brain regions, which could lead to different levels of signal-to-noise ratio for extracted time series after averaging across all voxels within these parcels. We agreed with this possibility, but hold the idea that it is more important to extract signals from biological-valid and meaningful parcels instead of parcels with externally forced equal size. The latter may cause the issue that averaged neural signals are the combinations of two or more rather separate signals which associate with different (cognitive) functions.

For each of the 314 parcels of the whole-brain parcellation, we compared it with the mask of *overlap network*, *inhibitory control network*, and *memory retrieval network* and identified the mask in which the parcel shared the highest percentage of common voxels. The parcel was assigned to that category if the highest percentage is higher than 10%. If the highest percentage of common voxels is lower than 10%, the parcel was not assigned to any category. After this procedure, 110 out of the 314 parcels were assigned to one of the categories. Specifically, 71 parcels were considered as *memory-related regions*, 29 parcels were categorized as *control-related regions*, and 10 parcels were labeled as *overlap regions* in our following analysis.

### Extraction of time series from parcels

We additionally removed nuisance time series (cerebrospinal fluid (CSF) signals, white matter signals, motion, and event-related activity) using a method based on a projection on the orthogonal of the signal space (Friston et al., 1994; Lindquist et al., 2019). We generated confounding time series (CSF, white matter, the six rigid-body motion parameters (three translations and three rotations), and framewise displacement (FD)) for each run of each participant. Event-related activity time series were estimated by a finite impulse response (FIR) function. A recent study has shown that the removal of event-related activity based on FIR modeling is an important step for the preprocessing of time series during a task (Cole et al., 2019). The signal from each parcel was extracted and z-scored, and all nuisance time series were removed simultaneously using the *nilearn.signal.clean* function. All cleaned time series were shifted 3 TRs (4.5 s) to account for the HRF delay and then aligned with the task demand (i.e., retrieval or suppression) at that moment. We shifted the time series by 4.5s because we reasoned that the peak of the HRF is between 4-6 s from the triggering event. This TR-wise data shifting is a pretty standard practice in time-resolved decoding of fMRI data (https://brainhack-princeton.github.io/handbook/content_pages/05-02-mvpa.html<u>#</u>)

### The transition of neural states analysis

First, we characterized the transition of neural states at the group level. Extracted time series from each run of each participant were split according to the task instruction (i.e., memory retrieval or memory suppression) and concatenated. Second, two kinds of time series were further concatenated across five TNT runs within that participant (*except for one participant, only four complete TNT runs were included*). Third, time series were concatenated across all participants. Fourth, two time-series were averaged across all time points to represent the mean activity intensity for that parcel during retrieval or suppression.

To estimate the relative dominance of each parcel during two neural states (i.e., Think and No-Think), we ranked the mean activity intensity of each parcel (the highest activity was ranked first). We then calculated the changes in ranks when the task switched from Think to No-Think by subtracting the rank during Think from the rank during No-Think. The same analyses were conducted with raw signal intensity and Z-values. Results can be found in the **Figure S4**. The negative change suggested an increase in relative dominance, while the positive change represented the opposite. We calculated two neural indexes (“state transition index” and “state transition index Version 2 (V2)”) to quantify the transition of neural states at the individual level and associate this individual difference with the subsequent suppression-induced forgetting effect. The state transition index was calculated by adding up the averaged relative decreases (absolute values for negative values) in rank values of *memory-related regions,* and the averaged relative increase in rank values of all *control-related regions*. The calculation and results of “state transition index Version 2 (V2)” can be found in the *Supplemental Material- An alternative method to quantify individual differences in neural state transitions*. It is notable that although transition index and transition index V2 were calculated using different methods, they were based on the same set of data. Therefore, the data analyses of index V2 should not be regarded as independent analysis. To explore whether the neural state transition during the TNT relates to the suppression-induced forgetting effect, we correlated two state transition indices with the *objective suppression score* and *subjective suppression score.* Suppression scores were calculated based on memory performance during the final memory test after TNT.

### Neural states decoding analysis

Before the decoding analysis, we generated the labels of task demand for each time point within the trial based on its instruction (i.e., Think or No-Think). For example, if the trial is a Think trial, time points started from the presentation of memory cues of this trial to the presentation of memory cues of the next trial were labeled as “Think.” We performed the time-resolved multivariate decoding analysis based on the brain activity of all 110 regions and corresponding labels of task demand during each time point. This decoding analysis allowed us to generate the predicted label of task demand for each time point, thus revealing the fast dynamics of the neural state transition induced by the switch of task demand. Specifically, decoding analysis via the linear Support Vector Classification (SVC), the C-Support Vector Machine within the scikit-learn package (https://scikit-learn.org/stable/). We used default parameters of the function (regularization (C)=1, radial basis function kernel with degree=3). The classification of neural states was performed separately for each time point using a leave-one-run-out cross-validation approach within each participant. This procedure resulted in a decoded task demand for each time point of each participant. These predictions were evaluated by comparing these decoded task demands with actual task demand. To separate all types of correct and incorrect classification for the following analyses, we generated the confusion matrix for each participant. This confusion matrix contained the percentage of all four situations based on the task demand and if the prediction matches the task instruction (i.e., Think- Correct classification, Think- Incorrect classification, No-Think- Correct classification, and No-Think- Incorrect classification). We extracted all SVC discriminating weights assigned to the features during the participant-specific decoding and averaged them across all participants to generate the neural state-predictive pattern. The brain parcels with higher absolute values contributed more to decoding models.

To test for possible differences in neural representations of task demand induced by the task-switching, we performed the described decoding analyses for switch time points and non-switch time points separately. The switch time points were defined as the presentation time (2TRs; 3s) of the first memory cue after the switch of task demand. The two decoding analyses yielded decoding accuracies for switch time points and for non-switch time points for each participant. We compared these two types of decoding accuracies using the paired t-test. Less accurate decoding was described as the evidence for the weaker representation of the current task demand in the literature (Waskom et al., 2014; Loose et al., 2017). Also, because we only have two task demands, less accurate decoding reflects the unsuccessful transition from the previous demand to the current demand according to the instruction. After the general comparison between decoding accuracies between the *switch* and *non-switch* condition, we further analyzed them as the function of time (i.e., TR), and separately for Think and No-Think. This approach allowed us to explore the demand-specific neural dynamics of task demand representation within the different phases of one trial.

Next, we aimed to investigate the behavioral relevance of the mismatch moments (i.e., *incorrect classification*) between task demand and the underlying neural state. Because we were mainly interested in the switch-induced mismatch, we first restricted our analyses to these switch time points and then extended them to the non-switch time points as an exploratory analysis. For each participant, we averaged the trial-by-trial behavioral performance during the TNT task based on whether the actual task demand matches with decoded task demands. This yielded retrieval performance and suppression performance for *match* and *mismatch* conditions. Paired t-tests were performed to examine the effect of mismatch on the performance of memory retrieval and memory suppression separately. The performance calculations and comparisons described above were repeated for non-switch time points as well.

### Alternative classifier training procedures and effects on our results

During our main neural state decoding analysis, fMRI time points regardless of their specific phase (e.g., memory cue, response, or fixation…) and corresponding performance were all labeled as one of the two states (i.e., Think or No-Think) and used for classifier training and testing. We investigated the effects of applying two alternative classifier training procedures on our reported results. We described the procedures for alternative classifier training A and B below: (A) only time points during the memory cue phase were used for classifier training and testing; during classifier training, only the data from trials that participants reported successful retrieval and suppression were used for classifier training to further increase the specificity of the classifier. Successful retrieval trials were defined as trials with “often” or “always” retrieval reports and successful suppression trials were defined as trials with “never” or “sometimes” intrusion reports. (B) A subsampling balancing step was used to make sure that, for each participant, an equal number of data points from the “Think” and “No-Think” state was used for the classifier training; to prevent classifiers to use any sequential information during training, fMRI data (together with their state labels) were re-ordered to not reflect its original temporal sequence. For both procedures (A) and (B), after training, classifiers were used to generate the neural states (i.e., predicted state labels) only for time points during the memory cue phase. These labels were then analyzed to replicate three key findings of our decoding analysis: (1) higher-than-chance level neural state decoding accuracies; (2) less accurate decoding after switching compared to non-switch trials; (3) when predicted state labels (i.e., neural state) cannot match with the actual task demand, trial-by-trial behavioral performance was compromised.

### Relationship between head motion, neural state transitions, and behaviors

To explicitly assess how head motion could potentially affect our results, we derived a time point-by-timepoint measure of head motion, framewise displacement (FD) (Power et al., 2012), during the TNT task. FD is defined as the sum (in mm) of rotational and translational displacements from the current volume to the next volume. We aligned the time-series of FD with task structure and behaviors in a way similar to the analyses of the time series of fMRI signals but did not consider the HRF. The following contrasts were performed to compare head motion between conditions: (1) difference in FD between Think trials and No-Think trials; (2) difference in FD between correct neural state decoding and incorrect neural state decoding; (3) difference in FD between the switch and non-switch condition. Correlations analyses were performed between individual differences in head motion between Think and No-Think trials (i.e., FD_Think_-FD_No-Think_), *state transition index*, *objective/subjective suppression score*.

### Data and code availability

All research data of this study were uploaded to the Donders Repository ( https://data.donders.ru.nl/) and are publicly available. The project was named *Tracking the in- voluntary retrieval of unwanted memory in the human brain with functional MRI* in the Repository ( https://doi.org/10.34973/5afg-7r41). Some data, such as statistical maps and brain parcels of interest were shared via the Neurovault Repository https://identifiers.org/neurovault.collection:7731). Supplemental Material can be found in OSF (((https://osf.io/cq96h/).

Behavioral data were analyzed by *JASP* (https://jasp-stats.org/). For the term-based meta-analysis of neuroimaging studies, we used the *Neurosynth* (https://neurosynth.org/), and *BrainMap* (http://www.brainmap.org/). Preprocessing of neuroimaging data was performed by *FSL* (https://fsl.fmrib.ox.ac.uk/fsl/fslwiki), *ICA-AROMA* (https://github.com/maartenmennes/ICA-AROMA), and *fMRIPrep* (https://fmriprep.readthedocs.io/en/stable/). Python packages, including *Nilearn* (https://nilearn.github.io/), *Nistats* (https://nistats.github.io/), *Pandas* (https://pandas.pydata.org/), and *Nunpy* (https://numpy.org/) were used for the analyses of time series. Machine learning algorithms were based on *scikit-learn* (https://scikit-learn.org/) and implemented via *Nilearn* (https://nilearn.github.io/). *Anaconda* (https://www.anaconda.com/) Python 3.6 was used as the platform for all the programming and statistical analyses. Custom Python scripts were written to perform all analyses described based on the mentioned Python packages; all code is available from the authors upon request and will be released via our OSF repository (https://osf.io/cq96h/) upon publication.

