## Supplementary Material for "Dynamic transitions between neural states are associated with flexible task-switching during a memory task"

**This PDF file includes:**

Supplementary Text

Figure S1-S9

Table S1-S3

**Correspondence:**

Wei Liu

School of Psychology,

Central China Normal University (CCNU),

No. 152 Luoyu Road, Hongshan District, Wuhan 430079,

Hubei Province, Wuhan, China.

**1. Behavioral performance during the final memory test**

During the final memory test, each memory cue was presented again, and the participant was instructed to rate the confidence of their memory for this association, and then classified the category of the associated picture. We examined the effect of memory retrieval and suppression during the TNT on the subsequent subjective (confidence rating) and objective (if they selected the correct category) memory. Three kinds of associations (i.e. retrieval association, suppression association, and control association) did not differ in their objective recall accuracy (F [2,26] =0.524, p=0.595, η² =0.02; **Figure S2A**). Our experiment design may explain the lack of the main effect of modulation on objective memory at the group level: all associations underwent overnight consolidation, thus difficult to be modulated (Liu *et al.*, 2016; Liu *et al.*, 2020). Crucially, replicating the previous study (Levy and Anderson, 2012), we found the suppression-induced forgetting at the individual level, and this effect was determined by individual differences in the efficiency of suppression during the TNT task. Specifically, participants who were more effective in suppressing intrusions (more negative *intrusion slope score*) during the TNT phase were the ones who show larger suppression-induced forgetting effects (r=0.411, p=0.03; **Figure S2C**). Next, analyzing the subjective memory, we found a significant effect of modulation on subjective memory (F [2,26] =5.928, p=0.005, η² =0.186; **Figure S2B**). Participants reported higher confidence for retrieval associations compared to control associations (t=3.35, p _holm_=0.007) and a trend towards higher confidence compared to suppression associations (t=2.172, p _holm_=0.07). Finally, we asked if modulation affected retrieval speed indexed by the RT during the final test. Even though we did not find a significant main effect of modulation (F [2,26]=2.905, p=0.06, η² =0.03; **Figure S2D**), recall of *RETRIEVAL ASSOCIATIONS* was faster compared to the recall of *CONTROL ASSOCIATIONS* (t(26)=-2.486, p=0.02, Cohen’s d=-0.47).

**2. Additional analyses of Think-to-NoThink neural state transition**

To describe further the reconfiguration, we divided all ROIs into three groups (*increased group, stable group, and decreased group*) based on their relative changes in rank. When the task demand changed from Think to No-Think, 47.88% of the *memory-related regions* showed the top one-third decrease in relative rank values and therefore belonged to the decreased group. Another 39.43% of the *memory-related regions* did not change extensively during the transition (*stable group*), and 12.67% of the regions showed increases in their ranks (*increase group*). *Control-related regions* demonstrated the opposite neural changes: 75.86% of them belonged to the *increased group*, with 17.24% and 6.89% of their regions belonged to a *stable group* or *decreased group* separately. For *overlap regions*, 50% of them belonged to the *increased group*, 40% of them belonged to the stable group, and only one region belonged to the decreased group. We further looked at the proportion of *memory-related regions, control-related regions, and overlap regions* within *increased, decreased*, and *stable* groups separately. A chi-square test of independence was performed to examine the relations between their functions (i.e., *memory-related, control-related, or overlap*) and which change group they belong to. The relation between these variables was significant (*X*^2^ =41.38, p<0.001). *Control-related regions* were more likely to be allocated to the increased group, while *memory-related regions* were more likely to be assigned to the decreased group. Specifically, among the increased group with a total of 36 regions (around 33.3% of all 110 ROIs analyzed), 61% of the regions were *control-related regions* (expected percentage=26.4%, p<0.001, one-side binomial test). By contrast, within the decreased group, 92% of the regions were *memory-related regions* (expected percentage=64.5%, p<0.001, one-side binomial test). Taken together, we found that during Think condition, *memory-related regions* showed relatively high neural activity compared to *control-related regions* and *overlap regions*. When the task demand changed from Think to No-Think, *memory-related regions* showed decreases in their relative contribution, while *control-related regions* demonstrated an increase in their activity ranks. These patterns of changes were not only presented when we analyzed the rank of activity among ROIs (**Figure S4A**) but also existed when we analyzed their raw (**Figure S4B**) or Z score (**Figure S4C**) of activity intensity.

**3. An alternative method to quantify individual differences in neural state transitions**

In the main text, we presented significant correlations between the state transition index and *objective/subjective suppression effect* (See Figure 2C and D). However, statistical tests were just around a significant level of p=0.05. To further validate the relationship between neural state transitions during the TNT and the subsequent forgetting effect, we used an alternative method (i.e., state transition index Version2 (V2)) to quantify individual differences in neural state transitions and performed the correlation again. This method is based on the additional analysis of the Think-to-NoThink neural state transition above. For each participant, all 110 ROIs were divided into three groups (i.e., *increased group, stable group, and decreased group*) based on their relative changes in rank**.** The state transition index V2 was defined as the sum of the percentage of *memory-related nodes* within the *decreased group* and percentage of *control-related nodes* within the *increased group*. As shown in **Figure S5** (*right panels*), state transition index V2 positively associated with the individual differences in objective suppression score (r=0.43, p=0.02), and tended to associate with subjective suppression score (r=0.36, p=0.06).

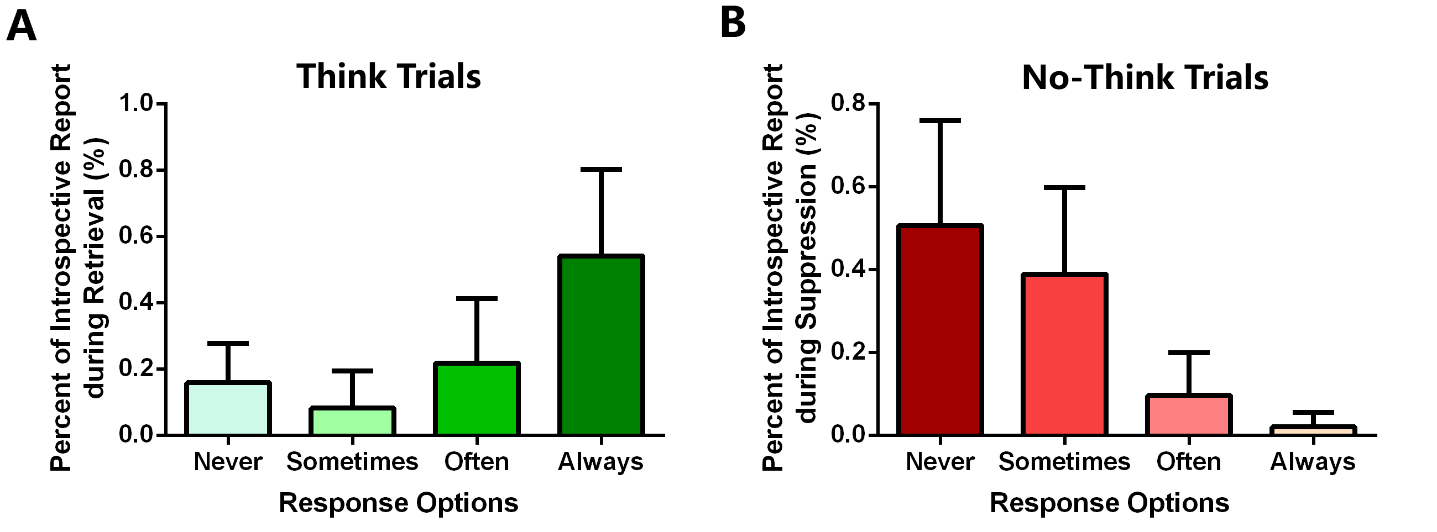

**Figure S1 Behavioral performance during the Think/No-Think task.** **(A)** Percentage of the trial-by-trial introspective report during the Think trials. For most of the Think trials, associated pictures were successfully recalled (1-P_never_: mean=84.05%, SD=11.79 %). **(B)** Percentage of the trial-by-trial introspective report during the No-Think trials. During half of the No-Think trials, participants successfully suppressed the tendency to recall the associated pictures (P_never_: mean=50.62%, SD=25.35%).

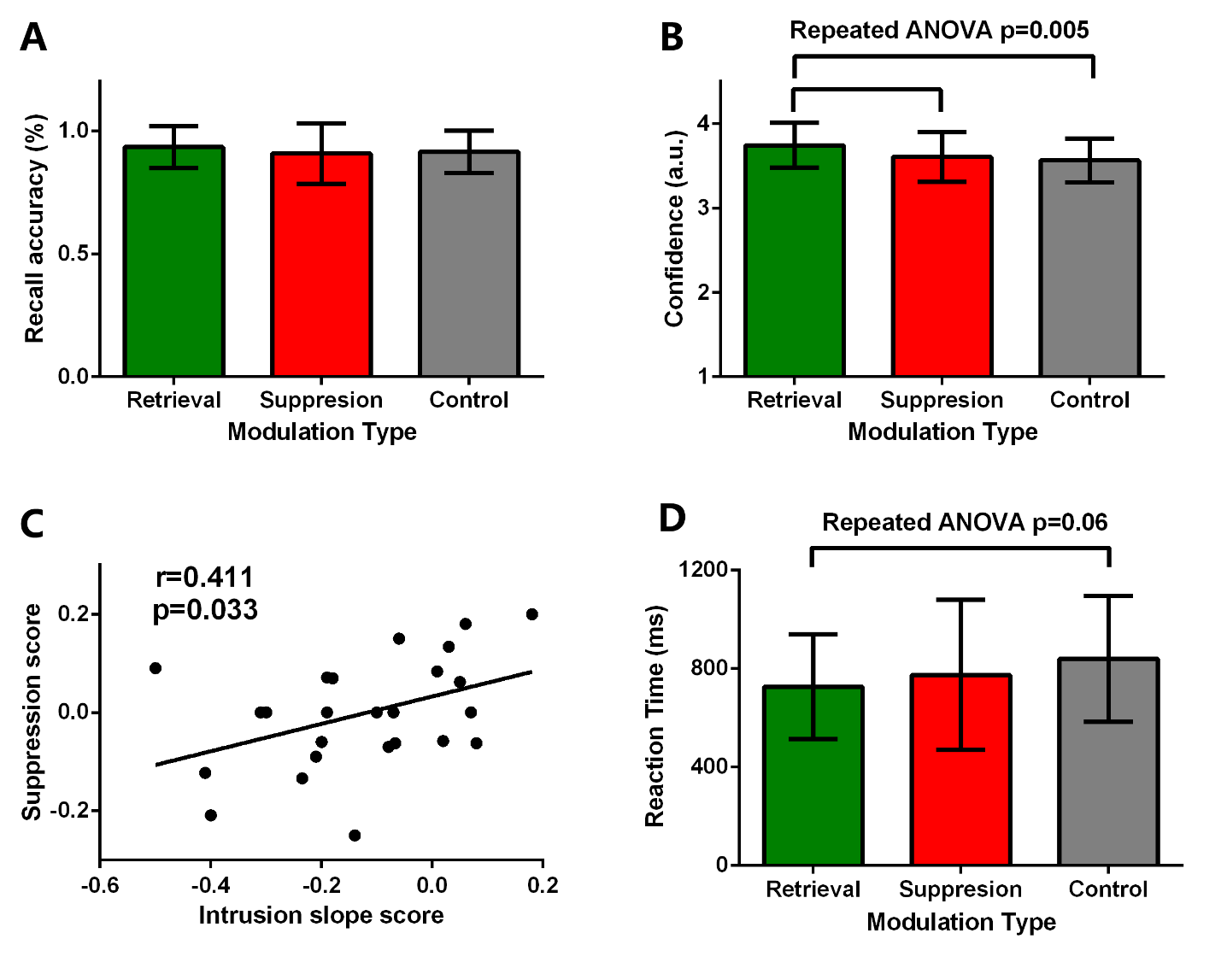

**Figure S2** **Behavioral performance during the final memory test. (A)** There is no effect of retrieval or suppression on the accuracy of the categorization during the final test (p=0.595). **(B)** For *RETRIEVAL ASSOCIATIONS*, participants reported higher subjective confidence compared to *SUPPRESSION ASSOCIATIONS* (t(26)=2.172, p_holm_=0.07, Cohen’s d=0.41) , and *CONTROL ASSOCIATIONS* (t(26)=3.35, P _holm_=0.007, Cohen’s d=0.64). **(C)** Participants who are more effective in reducing suppression failures (more negative the *Intrusion Slope Score*) were the ones who show more evidence suppression-induced forgetting (more negative the *Suppression Score*). **(D)** For *RETRIEVAL ASSOCIATIONS*, participants spent less time during categorization compared to the *CONTROL ASSOCIATIONS* (t(26)=-2.486, p=0.02, Cohen’s d=-0.47), and the effect between three conditions tend to be significant (F [2,26]=2.905, p=0.06, η² =0.03). a.u= arbitrary unit.

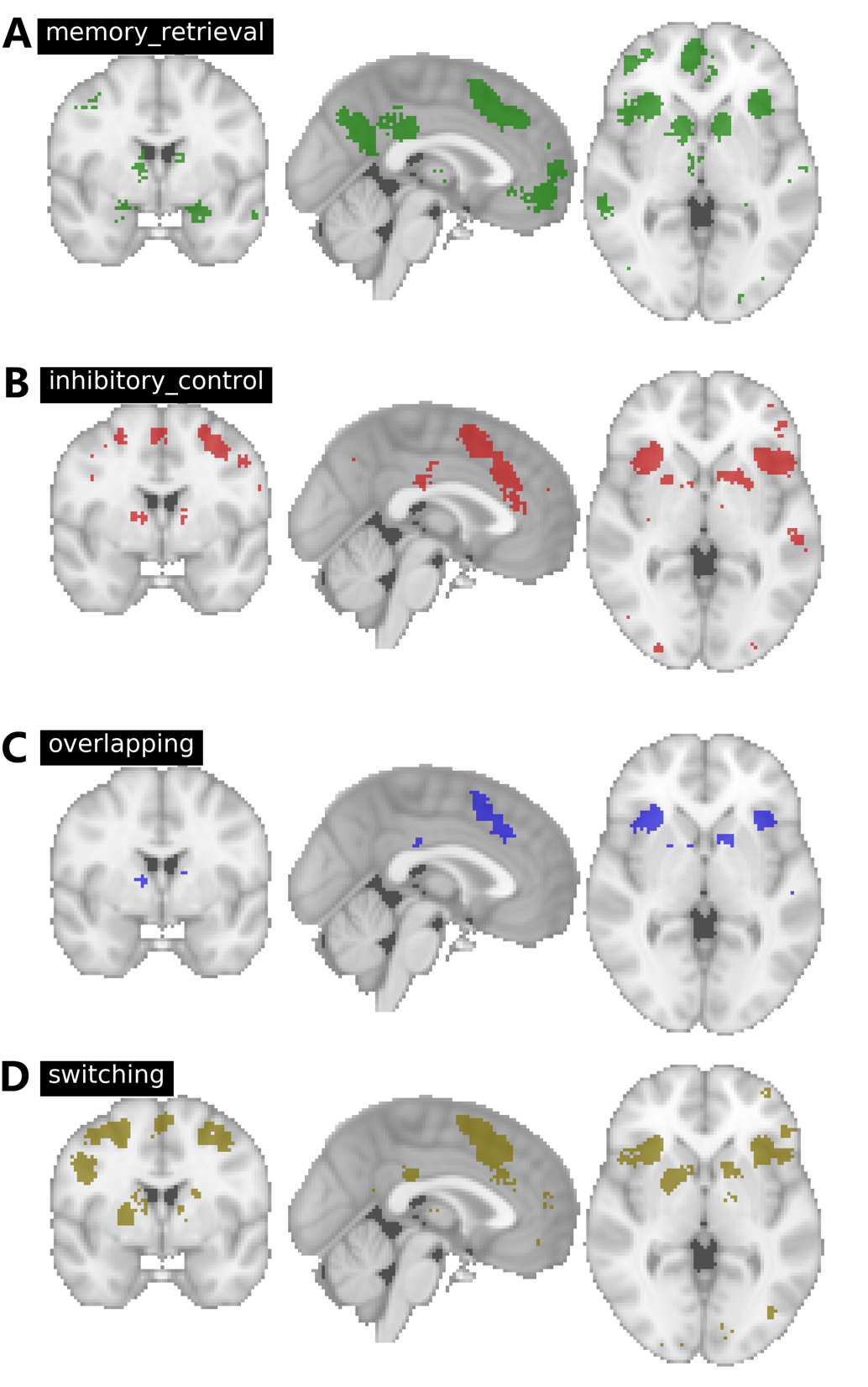

**Figure S3 Brain networks defined by the Neurosynth-based meta-analyses.** **(A)** Memory retrieval network generated using the term “memory retrieval.” **(B)** Inhibitory control network generated used the term “inhibitory control.” **(C)** Voxels that belong to both memory retrieval network and inhibitory control network. **(D)** Switching network generated used the term “task switching.” All raw statistical maps can be found in our Neurovault repository (<https://identifiers.org/neurovault.collection:7731>).

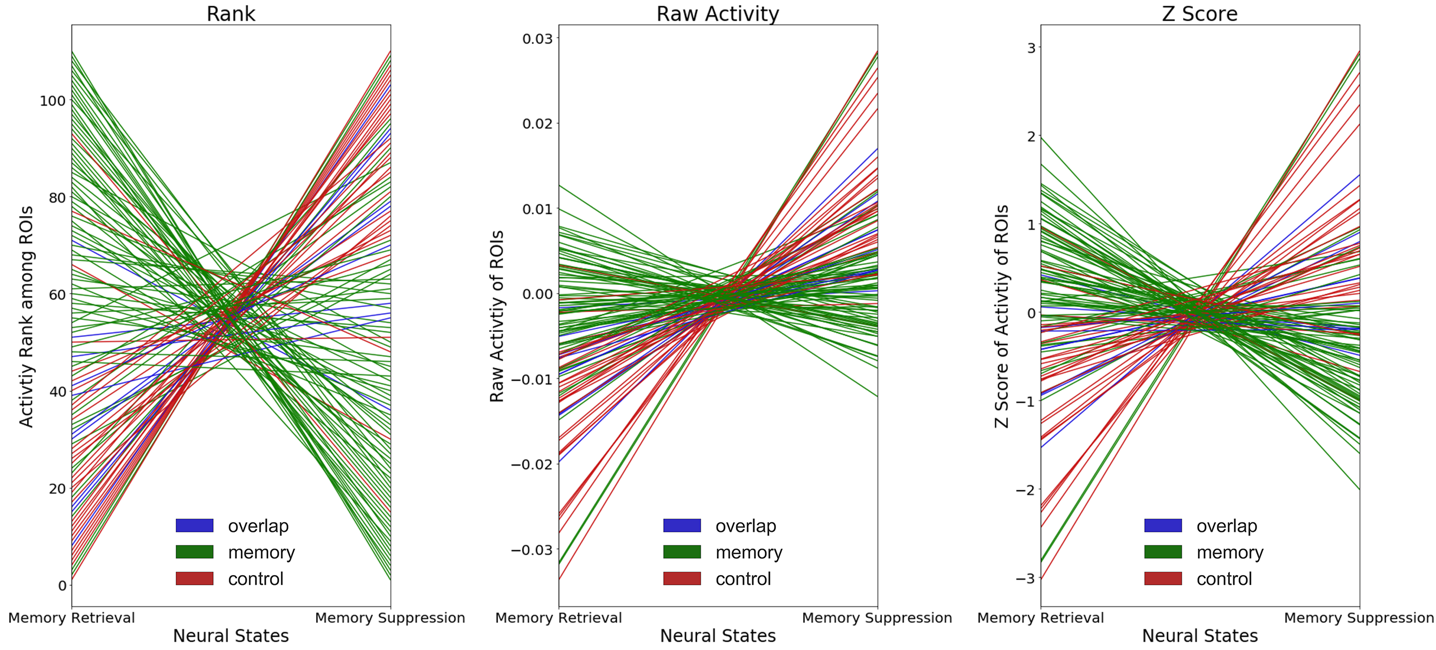

**Figure S4. Neural state reconfiguration during the Think-to-NoThink transition**. (A) Visualized based on the rank of activity intensity among ROIs. **(B)** Visualized based on raw activity intensities of ROIs. **(C)** Visualized based on the Z score of activity intensity of ROIs.

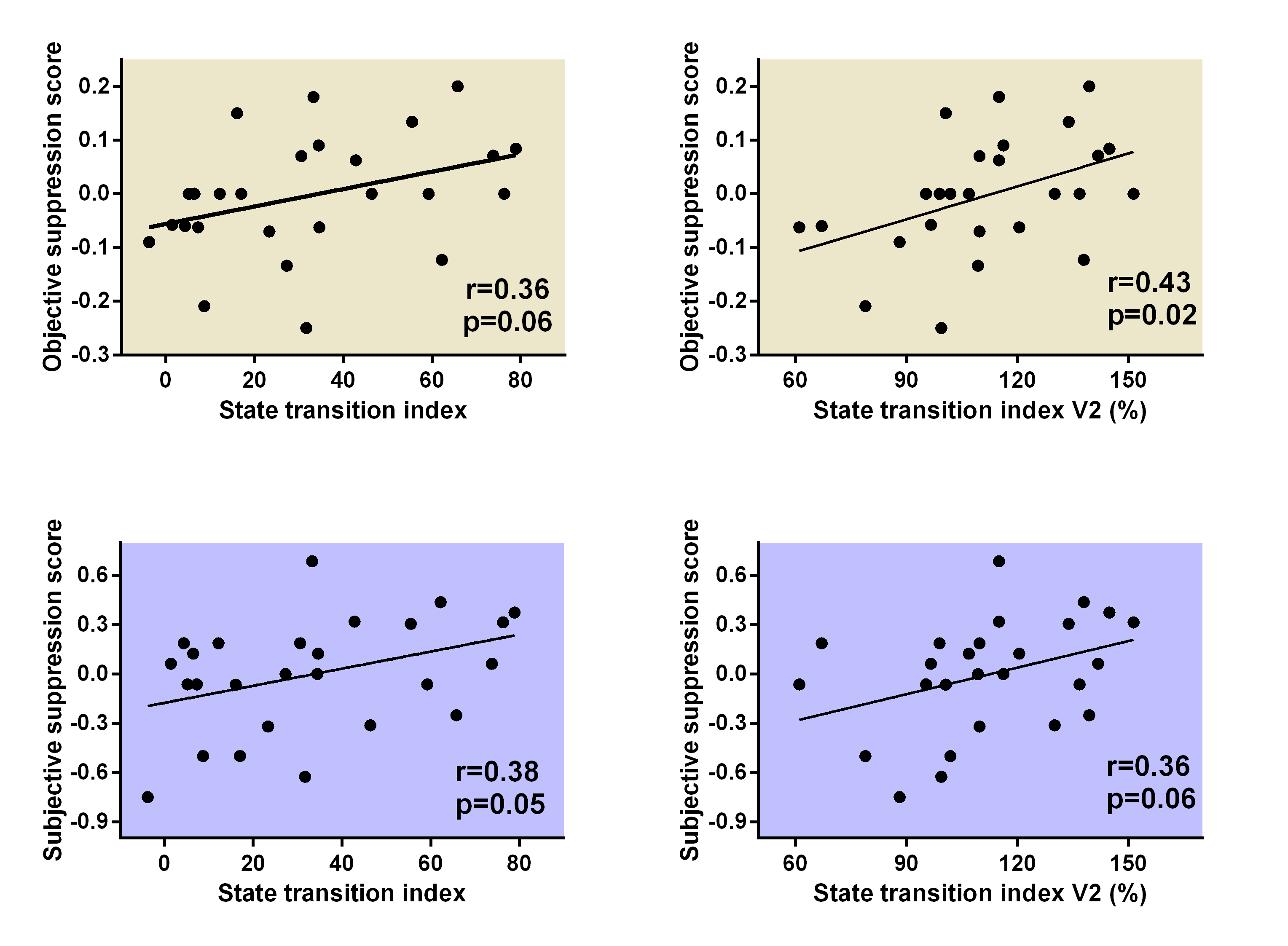

**Figure S5 Individual differences in two kinds of state transition index are correlated with both objective and subjective suppression score**. The results of the state transition index (i.e., left two panels) were presented in the *main text*. The results of state transition index V2 (i.e., right two panels) were presented in the *Supplemental Text Section3*.

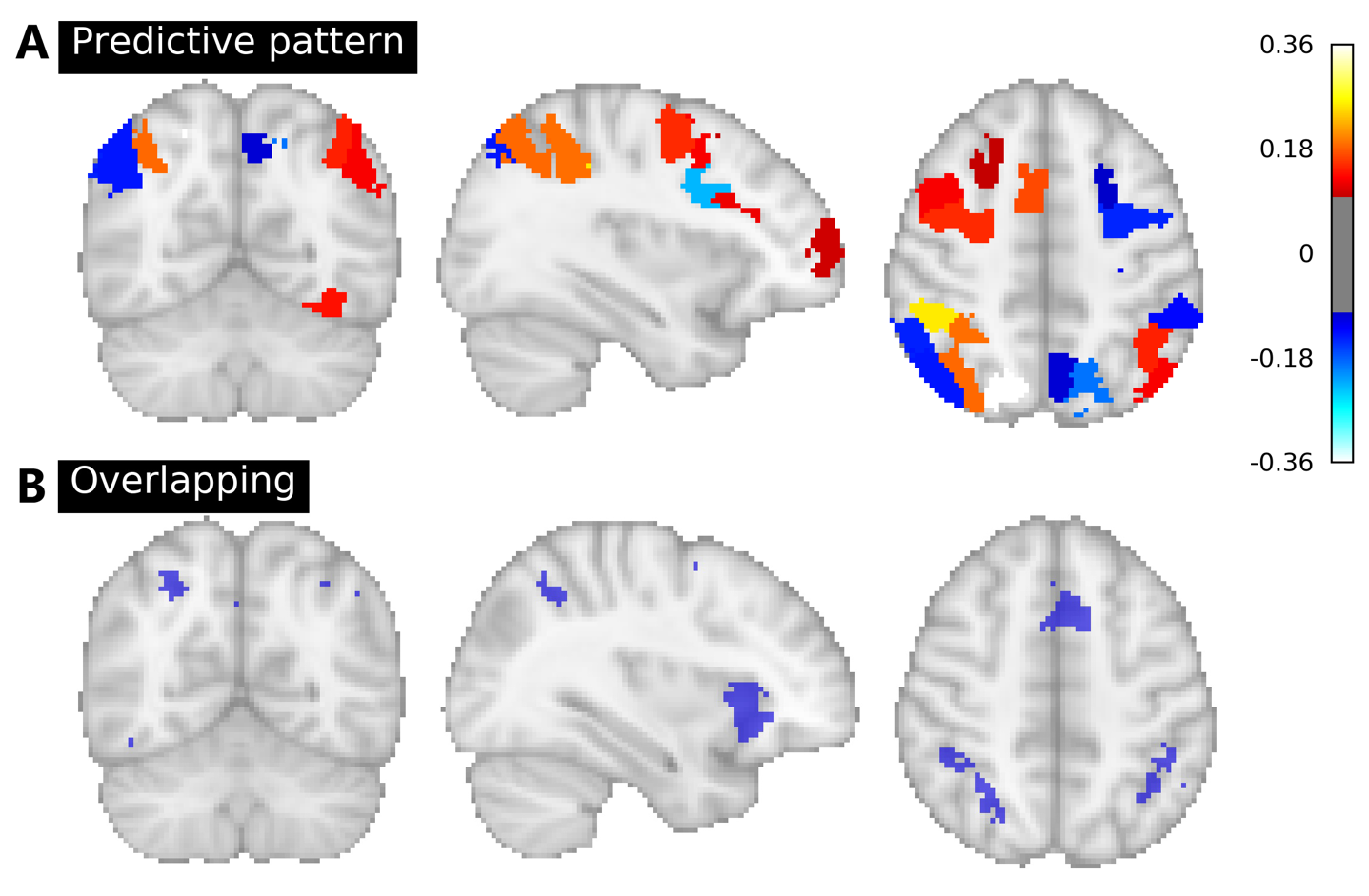

**Figure S6 Neural state-predictive pattern and overlapping network share similar spatial patterns.** **(A)** The contribution of different brain regions during the decoding. The map was visualized using an arbitrary threshold of 0.10. Higher absolute values of voxelwise statistical results represent a larger contribution during decoding. **(B)** Voxels that belong to both memory retrieval network and inhibitory control network (Neurosynth-based).

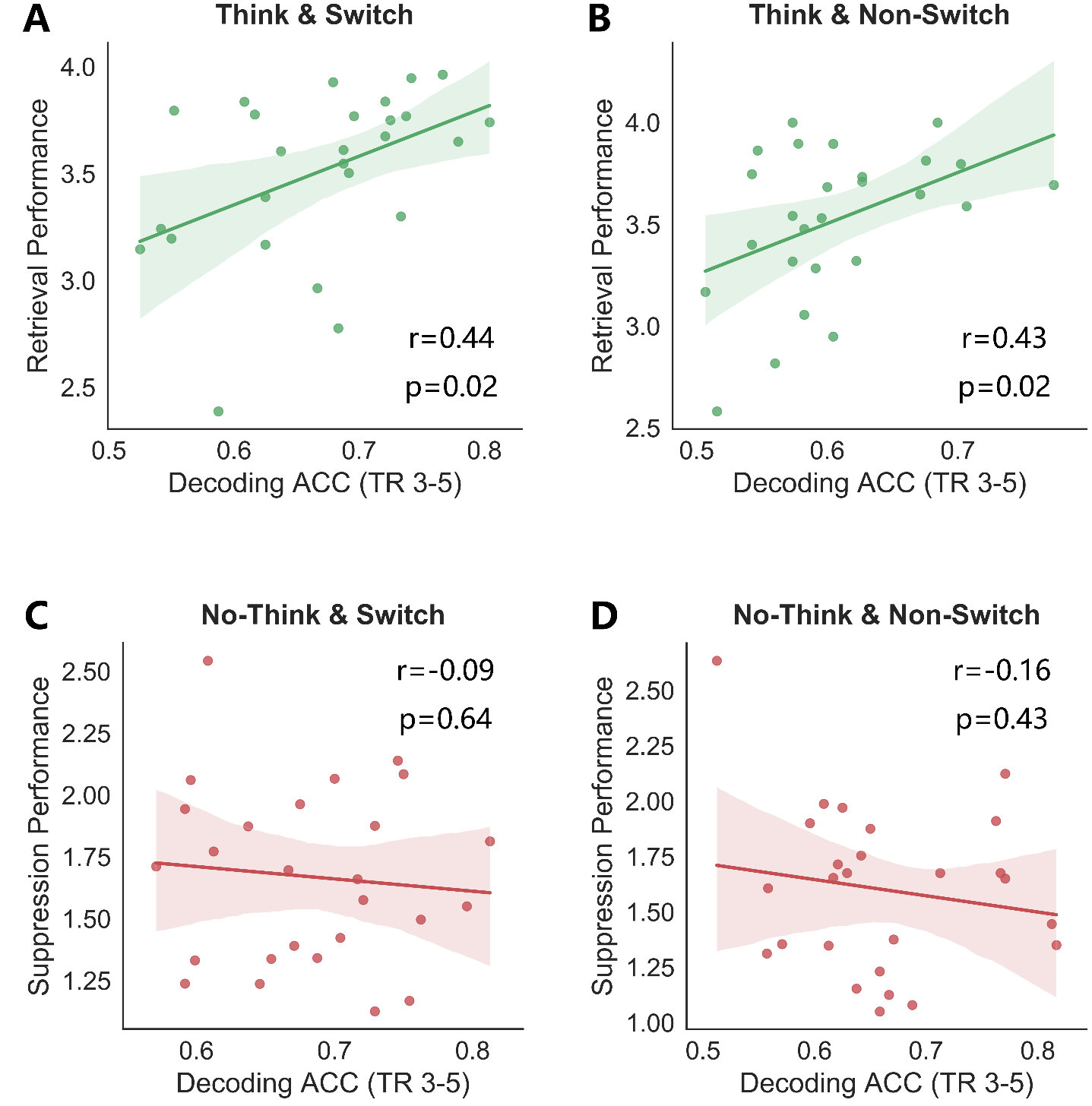

**Figure S7 Individual difference analyses between decoding accuracies (ACC) and behavioral performances. (A)** Stronger “adaptation” effects (*i.e., higher decoding ACC during the time window between TR=3 and TR=5*) correlated with better memory retrieval performance when it was a switch trial (r=0.44, p=0.02). **(B)** Stronger “adaptation” effects correlated with better memory retrieval performance when it was a non-switch trial (r=0.43, p=0.02). **(C)(D)** Decoding ACC during the same time window did not associate with memory suppression performance during either switch (p=0.64) or non-switch trial (p=0.43)

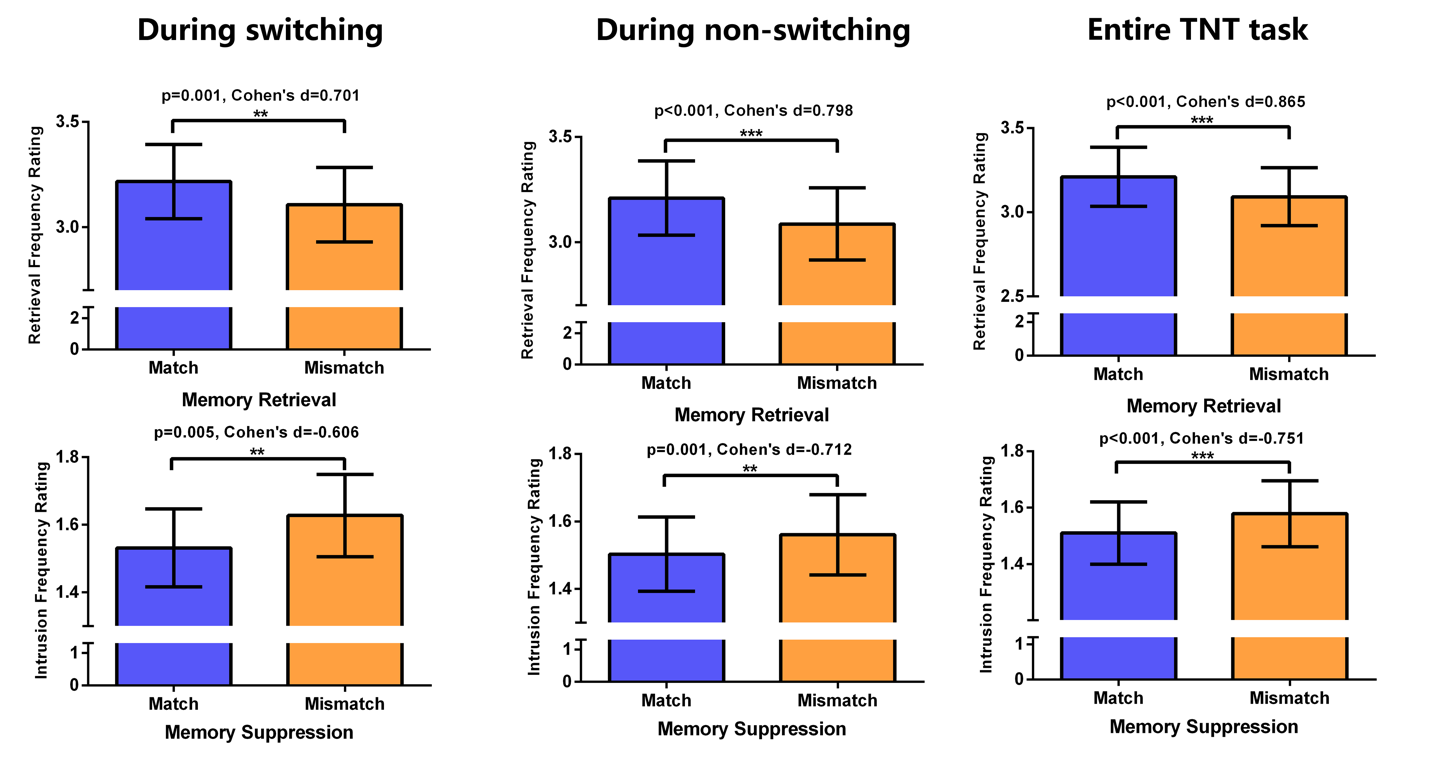

**Figure S8 Behavioral consequences of the mismatch between neural state and current task demand.** When the decoded neural state did not match with the task demand (i.e., Think decoded as No-Think), participants reported worse memory retrieval performance during Think trials. When the neural decoder misclassified No-Think moments as Think, participants reported more memory intrusions during No-Think trials. These two effects can be detected during the *switching* period (i.e., left panel), *non-switching* period (i.e., middle panel), and all time points within the *entire Think/No-Think* task (i.e., right panel)

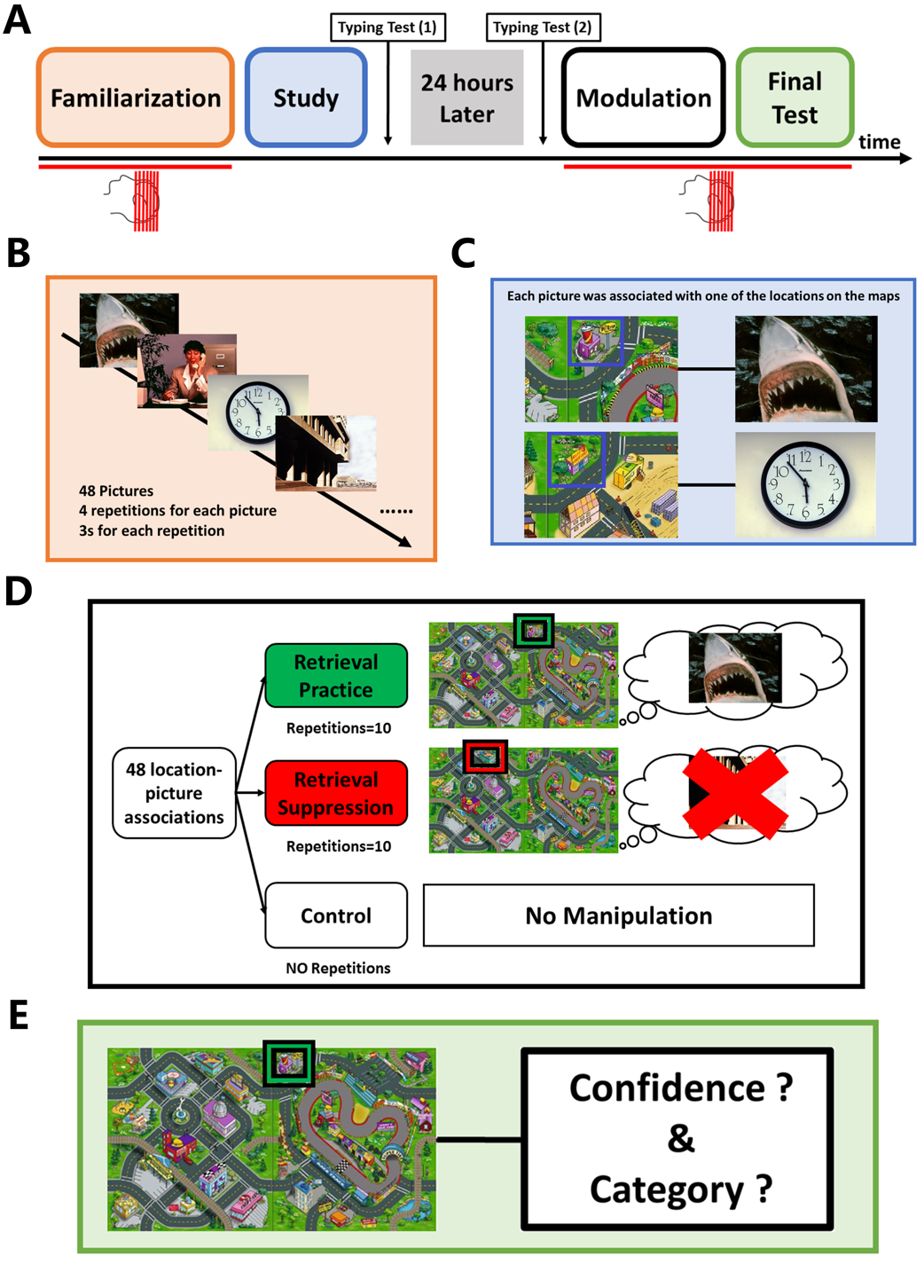

**Figure S9 Schematic of the experiment design. (A)** Timeline of the two-day experimental procedures. Red lines below the timeline indicate the tasks in the MRI scanner. **(B)** During the familiarization phase, all of the pictures of the to-be-remembered associations were randomly presented four times for the familiarization and estimation of picture-specific activation patterns. To keep participants focused, on each trial, they were instructed to categorize the picture shown as an animal, human, location, or object. **(C)** Study phase. Participants were trained to associate memory cues with presented pictures. **(D)** Modulation phase. After 24 hours, we used the Think/No-Think paradigm to modulate consolidated associative memories. Participants were instructed to actively retrieve associated pictures in mind (“retrieval”) or suppress the tendency to recall them (“suppression”) according to the colors of the frames (GREEN: retrieval; RED: suppression) around locations. **(E)** Final memory test phase. Participants performed the final memory test after the modulation. For each of the 48 location-picture associations, locations were presented again, and participants were instructed to report the memory confidence and categorize the picture that came to mind.

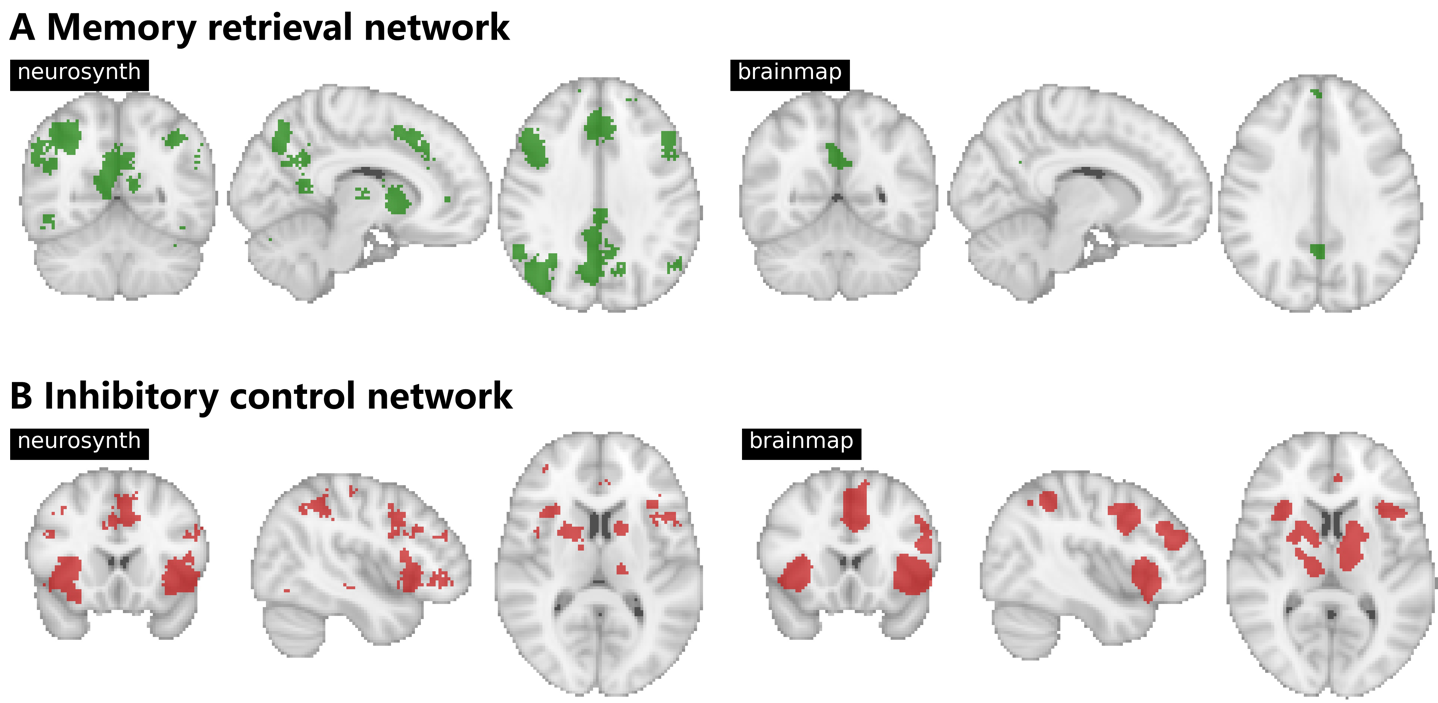

**Figure S10 Comparison between brain networks generated by the Neurosynth and Brainmap. (A)** Memory retrieval network. The network generated by the Neurosynth using the term “*memory retrieval*” (left panel). The network generated by the Brainmap based on activation-based mapping in healthy participants who perform the paradigm of “*episodic recall*” (right panel). **(B)** Inhibitory control network. The network generated by the Neurosynth using the term “*inhibitory control*” (left panel). The network generated by the Brainmap based on activation-based mapping in healthy participants who perform the paradigm of “*go/no-go*” (right panel)

**Table S1** **Significant activated clusters during No-Think trials compared to Think trials**

| **Contrast** | **Brain region** | **Hemisphere** | | **MNI coordinates** | **Cluster size** | **Peak voxel Value** |
| --- | --- | --- | --- | --- | --- | --- |
| **No-Think**  **>**  **Think** | IFG/Insula | | L | -38 22 8 | 920 | 5.5 |
|  | IFG/Insula | | R | 46 18 12 | 2397 | 5.5 |
|  | DLPFC | | R | 22 46 24 | 1681 | 4.5 |
|  | DLPFC | | L | -22 44 18 | 160 | 4.5 |
|  | IPL | | R | 56 -42 34 | 1305 | 5.5 |
|  | IPL | | L | -58 -54 42 | 226 | 4.4 |
|  | Thalamus | | R | 16 -22 2 | 677 | 6.6 |
|  | Precuneus | | R/L | 14 -60 56 | NA | 5.4 |
|  | Postcentral gyrus | | R | 45 -20 55 | NA | 7.4 |
|  | SMA | | R/L | 8 0 53 | NA | 5.5 |
|  | dACC | | R/L | 8 20 37 | NA | 4.4 |

IFG=Inferior Frontal Gyrus; DLPFC=Dorsolateral Prefrontal Cortex; IPL=Inferior Parietal Lobule; SMA=Supplementary Motor Area; dACC= dorsal Anterior Cingulate Cortex

**Table S2 Significant activated clusters during Think trials compared to No-Think trials**

| **Contrast** | **Brain region** | **Hemisphere** | **MNI coordinates** | **Cluster size** | **Peak voxel Value** |
| --- | --- | --- | --- | --- | --- |
| **Think**  **>**  **No-Think** | mPFC | R/L | 0 -44 -10 | 6573 | 4.03 |
|  | Insula | L | -42 -16 18 |  | 7.49 |
|  | ITG | L | -42 -42 -10 | 178 | 4.54 |
|  | STG/IPL/Precuneus | R | 54 -30 10 | 2925 | 5.36 |
|  | Hippocampus | R | 30 -40 4 |  | 3.74 |
|  | Precunues/PCC/SMA | L | -4 -40 34 | 6220 | 4.85 |
|  | Precentral/Postcentral Gyrus | L | -40 -36 60 |  | 7.32 |
|  | Posterior Cerebellum | R | 22 -60 -46 | 169 | 5.30 |
|  | Anterior Cerebellum | R | 16 -54 -20 | 1065 | 7.28 |

mPFC= medial Prefrontal Cortex; ITG= Inferior Temporal Gyrus; STG= Superior Temporal Gyrus; PCC= Posterior Cingulate Cortex; SMA= Supplementary Motor Area

**Table S3 Top 10 brain parcels with high classification weights during the neural state prediction**

| **Anatomical Label** | **X** | **Y** | **Z** | **Hemisphere** | **Network** | **Classification Weight** |
| --- | --- | --- | --- | --- | --- | --- |
| **SPL** | -34 | -48 | 46 | LH | DorsAttn | 0.189181458 |
| **LOC** | -16 | -72 | 54 | LH | DorsAttn | -0.358755616 |
| **Middle frontal gyrus** | -32 | -4 | 52 | LH | DorsAttn | 0.151391694 |
| **dACC** | -6 | 10 | 40 | LH | SalVentAttn | 0.169217997 |
| **IPL** | -34 | -66 | 48 | LH | Cont | 0.1866069 |
| **IPL** | -44 | -42 | 46 | LH | Cont | 0.255550192 |
| **Middle frontal gyrus** | -42 | 12 | 34 | LH | Cont | -0.229290806 |
| **IFG** | -54 | 20 | 12 | LH | Default | -0.179694589 |
| **SPL** | 16 | -72 | 54 | RH | DorsAttn | -0.190727893 |
| **Putamen** | NA | NA | NA | RH | Subcortical | -0.161437796 |

DorsAttn=dorsal attention network; SalVentAttn=salience ventral attention network; Cont=frontal parietal control network; Default=default network; SPL=superior parietal lobule; LOC= lateral occipital cortex; dACC= dorsal Anterior Cingulate Cortex; IPL= Inferior Parietal Lobule; IFG=Inferior Frontal Gyrus
